# Long-term live imaging of epithelial organoids and corresponding multiscale analysis reveal high heterogeneity and identify core regulatory principles

**DOI:** 10.1101/2020.07.12.199463

**Authors:** Lotta Hof, Till Moreth, Michael Koch, Tim Liebisch, Marina Kurtz, Julia Tarnick, Susanna M. Lissek, Monique M.A. Verstegen, Luc J.W. van der Laan, Meritxell Huch, Franziska Matthäus, Ernst H.K. Stelzer, Francesco Pampaloni

**Affiliations:** Physical Biology Group, Buchmann Institute for Molecular Life Sciences (BMLS), Goethe-Universität Frankfurt am Main, Frankfurt am Main, Germany; Faculty of Biological Sciences & Frankfurt Institute for Advanced Studies (FIAS), Goethe-Universität Frankfurt am Main, Frankfurt am Main, Germany; Department of Physics & Frankfurt Institute for Advanced Studies (FIAS), Goethe-Universität Frankfurt am Main, Frankfurt am Main, Germany; Deanery of Biomedical Science, University of Edinburgh, Edinburgh, United Kingdom; Experimental Medicine and Therapy Research, University of Regensburg, Regensburg, Germany; Department of Surgery, Erasmus MC – University Medical Center, Rotterdam, The Netherlands; The Wellcome Trust/CRUK Gurdon Institute, University of Cambridge, Cambridge, United Kingdom

## Abstract

Organoids are morphologically heterogeneous three-dimensional cell culture systems. To understand the cell organisation principles of their morphogenesis, we imaged hundreds of pancreas and liver organoids in parallel using light sheet and bright field microscopy for up to seven days. We quantified organoid behaviour at single-cell (microscale), individual-organoid (mesoscale), and entire-culture (macroscale) levels. At single-cell resolution, we monitored formation, monolayer polarisation and degeneration, and identified diverse behaviours, including lumen expansion and decline (size oscillation), migration, rotation and multi-organoid fusion. Detailed individual organoid quantifications lead to a mechanical 3D agent-based model. A derived scaling law and simulations support the hypotheses that size oscillations depend on organoid properties and cell division dynamics, which is confirmed by bright field macroscale analyses of entire cultures. Our multiscale analysis provides a systematic picture of the diversity of cell organisation in organoids by identifying and quantifying core regulatory principles of organoid morphogenesis.

**Graphical Abstract:** 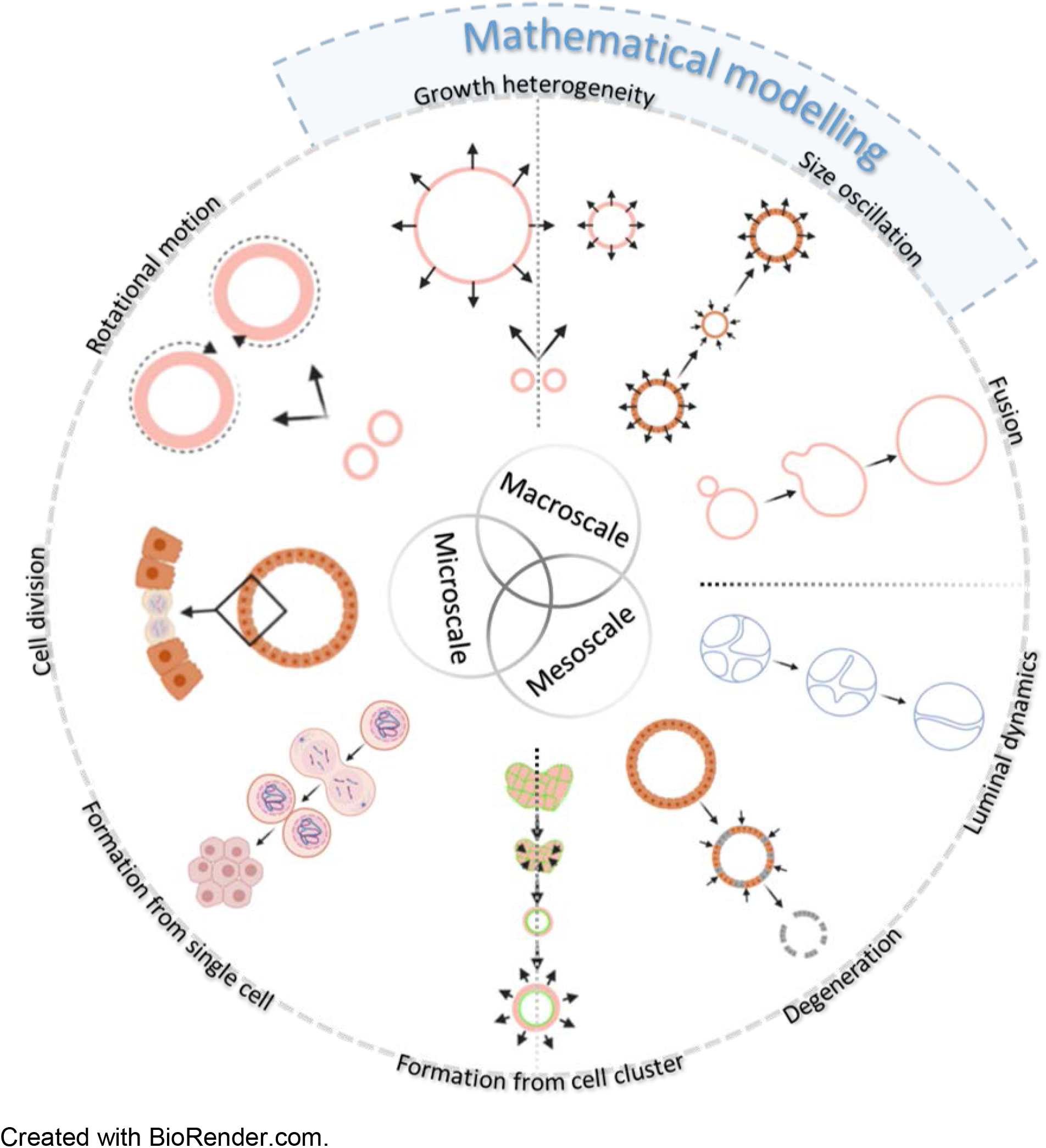

Created with BioRender.com.

## Introduction

Understanding the principles of collective cell behaviour in mammalian organs during development, homeostasis, regeneration and pathogenesis requires simplified models mimicking the *in vivo* cell-cell and cell-matrix interactions. To this aim, organoids provide an ideal *in vitro* model. Organoids are three-dimensional (3D) cultures obtained from pluripotent stem cells (embryonic or induced pluripotent stem cells) or organ-derived adult stem cells (Clevers, 2016). Organoid systems recapitulating the brain and the majority of epithelial organs have been established. These systems reproduce aspects of organ-specific development and disease (Kretzschmar et al., 2016; Lancaster et al., 2019) and are valuable for personalised (Broutier et al., 2017; de Winter-De Groot et al., 2018) and regenerative medicine (Takeda et al., 2013). Multicellular self-organisation determines organoid behaviour and morphology. For instance, epithelial organoids can acquire a spherical (“monocystic” (**Box 1**)), budding (“branched”), but also a dense (“polycystic” (**Box 1**)) phenotype (Loomans et al., 2018; Serra et al., 2019). Organoids are therefore a valid model to understand the principles of tissue self-organisation at the mesoscale, which are largely unknown (Trepat et al., 2018).

In order to fill this knowledge gap, a quantitative analysis at single-cell resolution is essential. A multiscale approach is required, capturing the cell-to-cell variability while monitoring the entire organoid system (Xavier da Silveira dos Santos et al., 2019) (**Box 1**).

The advancement of molecular biology allows quantifications of large-scale omics data at single-cell resolution. For example, high-throughput single-cell transcriptomics detect rare cell populations and trajectories between distinct cell lineages (Grün et al., 2015). Unlike most single-cell molecular characterisations, time-resolved advanced microscopy enables both spatio-temporal analysis of the organoids’ global morphology as well as “zooming-in” on the fates of a single cell. In previous studies, Bolhaquiero *et al.* (Bolhaqueiro et al., 2019) were able to combine single-cell molecular and image-based analyses and proved chromosome segregation errors with up to 18 hours long image acquisitions in a confocal microscope. In an approach using an inverted light sheet microscope, Serra *et al.* (Serra et al., 2019) were able to perform 5-day long live acquisitions of individual organoids.

Ultimately, the experimental quantitative data on organoid dynamics should serve as a foundation for mathematical models, which predict the experimental outcome and test hypotheses about underlying mechanisms of observed behaviours by altering controllable parameters *in silico* (Sasai, 2013).

In our study, we focus on two types of organoids initiated from adult progenitor cells of the pancreas and liver tumour as representatives for a spherical as well as a polycystic phenotype. Murine pancreas-derived organoids (mPOs), are used as a model to study pancreas development and the regeneration of pancreatic β-cells (Huch *et al.*, 2013). Human cholangiocarcinoma-derived organoids (hCCOs), are promising models to study personalised treatment of primary liver cancer (Broutier et al., 2017). So far, the morphology of these organoids was analysed qualitatively by immuno-fluorescent marker localisation and bright field observations at single time points.

Here, in order to assess cellular dynamics in organoid cultures and identify their morphological organisational principles, we developed two complementary image-based analysis pipelines covering multiple scales: (1) A “Light sheet pipeline”, based on light sheet-based fluorescence microscopy (LSFM), addressing the micro- (single cell) and the mesoscale (individual organoid). (2) A “Bright field pipeline”, based on bright field microscopy, accessing both the meso- and macroscale (entire organoid culture) (**Supplementary Figure 1**).

The light sheet pipeline relies on image acquisition with the Lightsheet Z.1 (Zeiss) microscope, for which we further developed a custom FEP-foil cuvette (Hötte et al., 2019) (*further referred to as Z1-FEP-cuvette*, **Box 1**) as sample holder. Based on tagging cells with fluorescent fusion-proteins, this setup allowed for observations of cellular dynamics in live epithelial organoid cultures for up to seven days while retaining optimal physiological conditions (3D ECM environment, precisely defined medium, constant pH, and controlled temperature). Primary cell cultures, such as organoids, require minimal exposure to phototoxic effects, which is given by low energy exposure, due to a fast image acquisition as well as z-plane-confined illumination in LSFM (Keller et al., 2008; Stelzer, 2015). The confined illumination also yields a higher axial resolution compared to other microscopy systems, while still facilitating the acquisition of a large samples (Greger et al., 2007; Verveer et al., 2007). These features enable us to monitor large numbers of organoids simultaneously (approx. 100-200 organoids, depending on seeding density) in a maximum volume of about 8 mm^3^. Next to the detailed visualisation of highly dynamic processes during organoid formation, the data acquired by LSFM allow for tracking, extraction and quantification of several organoid features, including cell number and organoid volume at the micro- and mesoscale.

Based on the quantitative data obtained from the light sheet pipeline (cell count and cell division rates), we developed a 3D agent-based model (**Box 1**) to investigate the underlying mechanics driving single-organoid behaviour. Such models provide a technique to represent a wet-lab experiment under idealised conditions (Karolak et al., 2018). In contrast to the basic mechanisms proposed by the model of Ruiz-Herrero *et al.* (Ruiz-Herrero et al., 2017), which describes the dimensionless radius for hydraulically gated oscillations in spherical systems, we built a full elastic 3D model, based on the core principles formulated in Stichel *et al.* (Stichel et al., 2017). We expect organoids to have a similar morphogenic behaviour to other 3D-cell cultures, such as MDCK-cysts and epithelial tissues, namely, they grow by mitosis, display an apical-basal polarisation (Odenwald et al., 2017) and secrete osmotically active substances into their lumen (Ishiguro et al., 2012). Further, we assume that neighbouring cells are tightly connected via cell-cell junctions (Harris et al., 2010) and the cell layer ruptures if the internal pressure reaches a critical point.

The single-cell resolution achieved by the light sheet pipeline is necessary for studying single-cell dynamics and collective cell dynamics in individual organoids in depth. However, the large amounts of data acquired by this pipeline require considerable computational resources, which hinder the extraction and quantification of macroscale (entire organoid culture) features. We therefore developed the bright field pipeline that measures luminal size changes at individual-organoid resolution based on projected luminal areas (**Box 1**). This pipeline enables the observation of entire organoid cultures (approx. 100-200 organoids within 25 μl ECM droplets, depending on seeding density) over several days while retaining optimal physiological conditions. In addition, the bright field setup allows label-free image acquisition, which ensures minimal exposure to phototoxic effects. Quantification of the projected luminal areas over time yields features on a mesoscale level, such as minimal and maximal area of individual organoids, which are used to determine the median area increase of the entire culture at the macroscale level.

Our light sheet data indicate that epithelial organoids show size oscillations (expansion and decline phases) (**Box 1**), which are frequently observed in small organoids (diameter < 400 μm), but much less in large organoids (diameter > 400 μm). This is reflected in our 3D agent-based model, which indicates the size oscillations arise in response to an interplay of an increase of the internal pressure, the cell division dynamics and the mechanical properties of the single cells. The critical internal pressure due to release of osmotically active substances into the lumen is reached earlier in organoids with increased surface-to-volume ratios (small organoids) compared to organoids with reduced surface-to-volume ratios (large organoids). We further verified these findings by quantifying the size oscillations in entire organoid cultures using the bright field pipeline.

In summary, our approach reveals the dynamics of organoid cultures from single-cell and single-organoid scale to the complete culture scale, ascertaining the core regulatory principles (**Box 1**) of their multicellular behaviour.

## Results

### Long-term live imaging with LSFM allows detailed visualisation of dynamic processes in organoid morphogenesis and reveals high heterogeneity in single-cell and individual-organoid behaviour

To gain deeper insights into the dynamic cellular processes occurring within organoid systems, we developed Z1-FEP-cuvette holders for live imaging with the Zeiss Lightsheet Z.1 microscope system (**Supplementary Figure 2**). As previously described (Hötte et al., 2019), ultra-thin FEP-foil cuvettes are sample holders for LSFM which preserve physiological culture conditions for organoid cultures and allow the acquisition of high resolution images at single-cell level. Using the Z1-FEP-cuvette, we recorded the formation and development of hCCOs expressing H2B-eGFP (nuclei marker) and LifeAct-mCherry (F-actin cytoskeleton marker) and mPOs expressing Rosa26-nTnG (nuclei marker) for up to seven days. The medium was exchanged every 48 hours to ensure sufficient nutrient supply. Temperature and CO_2_ levels were controlled to ensure optimal growth conditions (**Supplementary Figures 3, 4**). The setup enabled us to monitor dynamic processes at high temporal and spatial resolution in up to 120 organoids simultaneously contained in one Z1-FEP-cuvette (in this example in a total volume of 5.2 mm^3^ of technically possible 8 mm^3^) (**Supplementary Figure 5a**; **Supplementary Movie 1**). The images acquired by LSFM allow for detailed qualitative inspections and detailed feature tracking of several dynamic cellular processes at single-cell resolution (**Supplementary Movie 2**).

Visual inspection of the acquired data revealed that the initially seeded organoid cell clusters contract before the cells within the clusters start to rearrange and form spherical structures (**Figure 1**, **Formation**). The cells within these spherical structures begin to polarise and form a lumen (in this example around 13.5 hours), indicated by a stronger F-actin signal at the apical (luminal) side of the cell membranes. Potentially dead cells accumulate within the lumen, indicated by loss of the LifeAct-mCherry signal and by smaller nuclei with stronger H2B-eGFP signals, hinting towards apoptotic nuclear condensation (Mandelkow et al., 2017). The polarisation of cells in the epithelial monolayer is maintained during luminal expansion and is still clearly visible at later stages of organoid development (in this example around 41.0 hours) (**Figure 1**, **Polarisation**). The recording interval of 30 minutes, also allows us to visually track single cell division events (here: over a time course of 2.5 hours) (**Figure 1**, **Cell division**). We were also able to observe polarisation and cell division events in isolated single cells (**Supplementary Figure 5b**), which remained dormant for relatively long periods during observation. We identified an overall shrinking of the organoid, nuclear condensation and a fading nuclei signal to be hallmarks of organoid degeneration (**Box 1**) (**Figure 1**, **Degeneration**; **Supplementary Movie 1**). This process is initiated upon extended culturing without further medium exchange (here: after about 100 hours).

**Figure 1:**
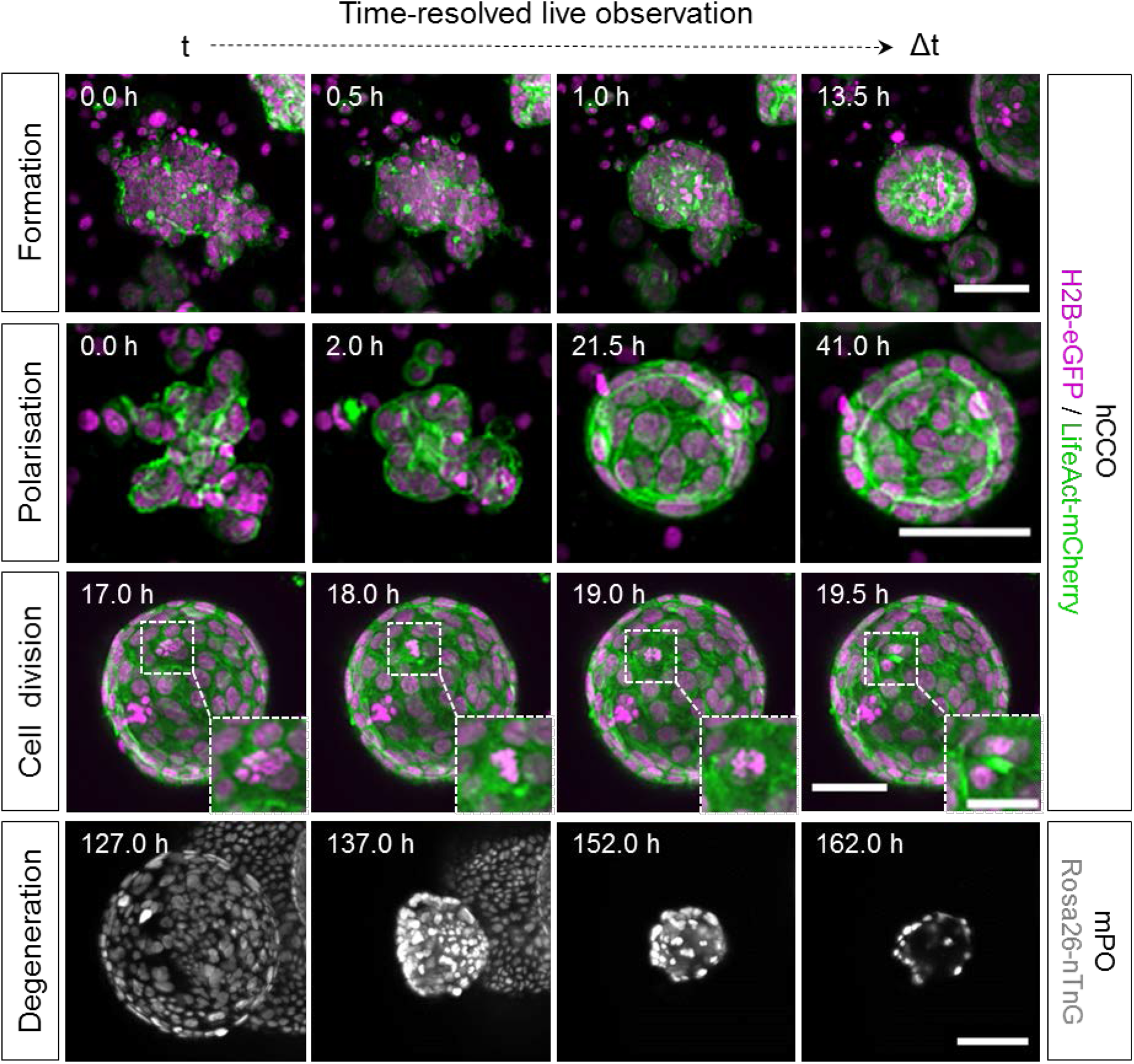
Time-resolved live LSFM recordings provide data for detailed qualitative inspections of dynamic morphological processes in organoid development. hCCOs and mPOs were seeded into Z1-FEP-cuvettes for long-term live observations. They expressed the nuclei marker H2B-eGFP (magenta) or Rosa26-nTnG (grey) and the F-actin cytoskeletal marker LifeAct-mCherry (green). About 120 organoids were recorded in image stacks up to 900 z-planes deep for at most seven days. The figure shows excerpts of maximum intensity z-projections. Microscope: Zeiss Lightsheet Z.1; objective lenses: detection: W Plan-Apochromat 20x/1.0, illumination: Zeiss LSFM 10x/0.2; laser lines: 488 nm, 561 nm; filters: laser block filter (LBF) 405/488/561; voxel size: 1.02 × 1.02 × 2.00 μm^3^; recording interval: 30 min; scale bars: 50 μm, 25 μm (inset).

Image-based segmentation and three-dimensional (3D) volume rendering of the acquired data allow for even more detailed inspections of features observed in the highly dynamic organoid system. While most processes can already be followed in maximum intensity z-projections of the acquired image data, 3D volume rendering facilitates a more detailed understanding of the underlying cellular dynamics from different perspectives. We identified organoid fusion to be a frequent phenomenon in the investigated cultures (**Figure 2**, **Fusion**; **Supplementary Movie 3**). After the epithelial monolayers of both organoids touch each other, they initiate an opening connecting both lumens. This opening then expands while cells migrate into one connected monolayer. Similar dynamics of cells migrating into one connected monolayer were observed in the formation, subsequent retraction and eventual rupture of duct-like structures within the lumen of a large organoid (diameter: ≥ 500 μm), which presumably emerged from fusion of multiple organoids (**Box 1**) (**Figure 2**, **Luminal dynamics**– ***lower panel***; **Supplementary Movies 4**, **5**). Volume rendering of cell nuclei revealed that small organoids (diameter at end of observation (48 hours): 100 μm) with large nuclei (longest axis: 52 μm) show less cell divisions and overall less cell movement than larger organoids (diameter at end of observation (48 hours): 180 μm) with smaller nuclei (longest axis: 32 μm) (**Supplementary Movie 6**). This further underline the heterogeneity in the investigated organoid systems.

**Figure 2:**
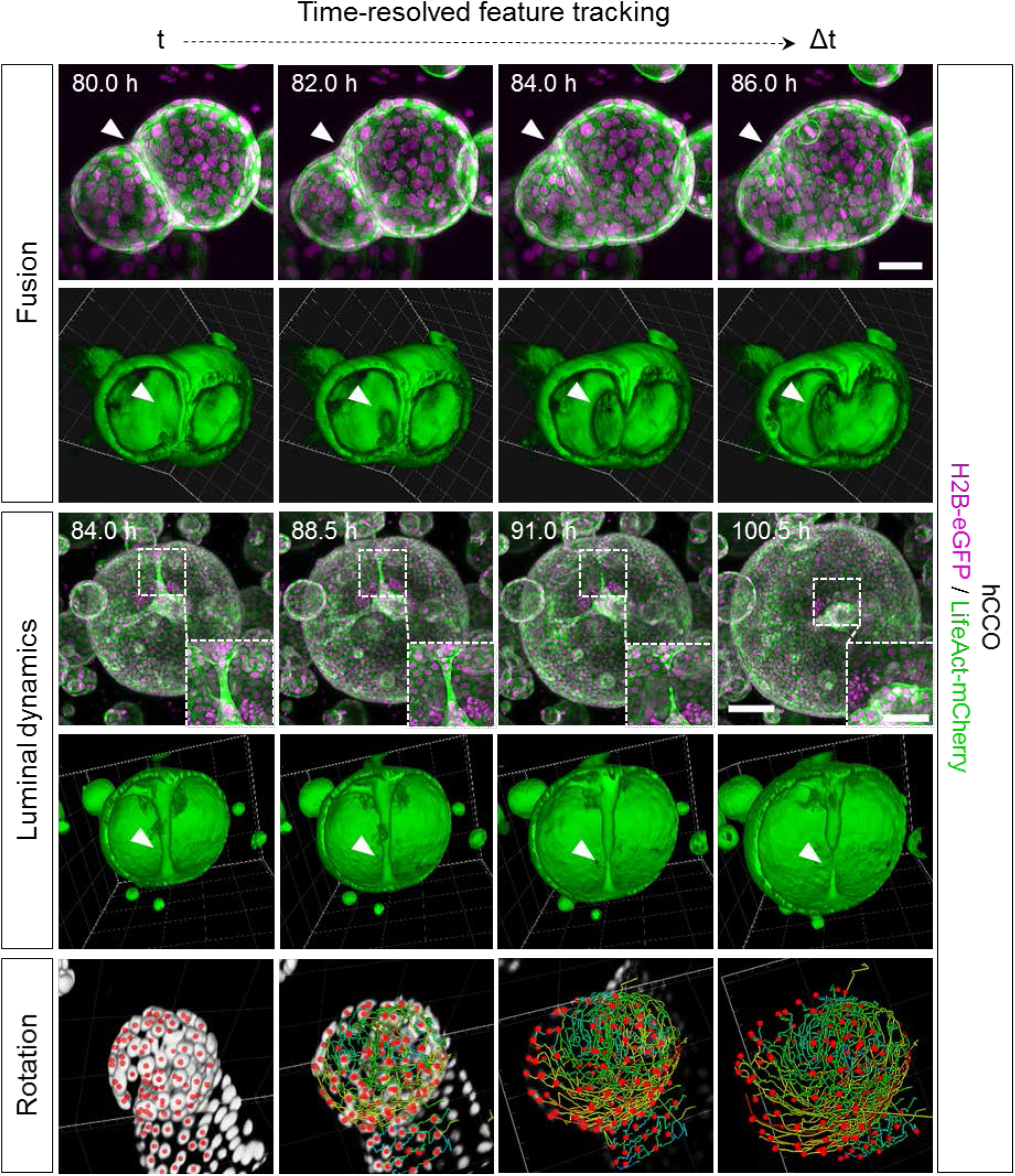
High-quality live LSFM image data provide an excellent basis for volume rendering and detailed feature tracking for the quantitative description of cellular dynamics in organoid development. 3D renderings offer detailed views into processes such as organoid fusion and elucidate the spatial context in observed luminal dynamics. 3D cell tracking reveals the complex rotation of the epithelial cell monolayer. hCCOs (seeded and maintained in Z1-FEP-cuvettes) expressed the nuclei marker H2B-eGFP (magenta) and the F-actin cytoskeletal marker LifeAct-mCherry (green). The figure shows segmented and tracked cell nuclei (Rotation; centroids – red; tracks – rainbow), excerpts of maximum intensity z-projections and 3D renderings of corresponding data sets. Segmentation, tracking and 3D rendering were performed with Arivis Vision4D. Microscope: Zeiss Lightsheet Z.1; objective lenses: detection: W Plan-Apochromat 20x/1.0, illumination: Zeiss LSFM 10x/0.2; laser lines: 488 nm, 561 nm; filters: laser block filter (LBF) 405/488/561; voxel size: 1.02 × 1.02 × 2.00 μm^3^; recording interval: 30 min; scale bars: Fusion −50 μm, Luminal dynamics – 100 μm, 50 μm (inset).

The observed heterogenic dynamic processes in organoids and organoid systems were quantitatively described using feature extraction and feature tracking tools. Single-cell tracking revealed that the previously observed cell movement in larger organoids with smaller cell nuclei can be described as a uniform rotation of the epithelial cell monolayer (**Figure 2**, **Rotation**; **Supplementary Movies 5**, **6**).

Single-cell tracking also revealed that the cells within an organoid move with different speeds during organoid expansion. Cells at the organoid’s “poles” tend to show less or slower movement (2 μm per hour) compared to cells located at the organoid’s equatorial plane (7 μm per hour) (**Supplementary Movie 7**). Furthermore, we observed that prior to organoid formation, some of the initially seeded cell clusters migrate through the ECM before they start to form a spherical structure (**Supplementary Movie 8**). In this example, the cell cluster travels at an average speed of 10 μm per hour (maximum speed: 23 μm per hour), covering a distance of about 250 μm in total.

### Long-term single-cell analysis of pancreas-derived organoids reveals cell-to-cell heterogeneity in cell proliferation

Next, we aimed for a deeper quantitative analysis of the dynamic cellular processes in luminal expansion. The collected high-resolution LSFM images, enabled the semi-automatic segmentation and quantitative feature extraction over the course of a 6 days acquisition. This provided robust, time-resolved data on cell nuclei numbers, organoid volume, surface area, the number of neighbouring cells for each cell as well as the cell density.

Using our previously published segmentation pipeline (Hötte et al., 2019; Schmitz et al., 2017), we processed one time-lapse dataset, which resulted in a total number of 288 segmented time points. From the segmented data, we chose to analyse three representative organoids. One small organoid (diameter < 400 μm), one large organoid (diameter > 400 μm) and one which was size-comparable to the large organoid but showed a higher cell number. The three mPOs expressed Rosa26-nTnG as a nuclei marker (**Figure 3**).

**Figure 3:**
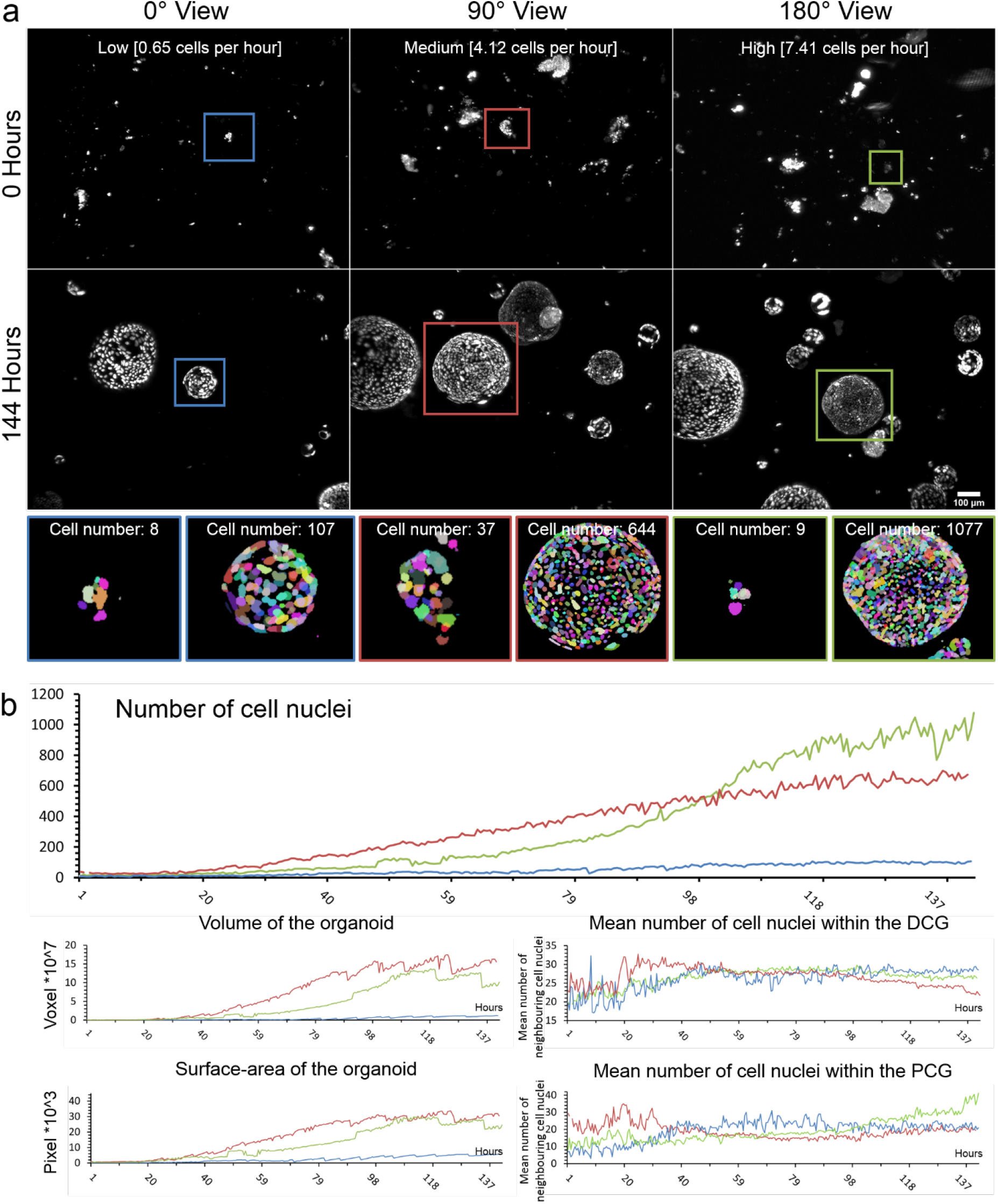
Long-term single-cell analysis of mPOs reveals heterogeneity of proliferation potentials. mPOs (seeded and maintained in one Z1-FEP-cuvettes) expressed the nuclei marker Rosa26-nTnG (grey). Organoids were imaged for six days and analysed with our previously published nuclei segmentation pipeline (Schmitz et al., 2017). (**a**) Three representative organoids are shown directly after seeding (0 h, upper row) and after six days (144 h, lower row). Every row shows one view of the same Z1-FEP-cuvette. Hence, all displayed organoids were grown simultaneously within one FEP-cuvette. The close-ups display the segmentation of the organoid at the corresponding time point. Different colours refer to individual cell nuclei. The coloured frames indicate organoids with different proliferation rates - green/high, blue/low and red/medium. (**b**) From top to bottom, corresponding evaluations of volume, surface area and neighbourhood relationships (DCG: Delaunay cell graph; PCG: proximity cell graph). Microscope: Zeiss Lightsheet Z.1; objective lenses: detection: W Plan-Apochromat 20x/1.0, illumination: Zeiss LSFM 10x/0.2; laser lines: 561 nm; filters: laser block filter (LBF) 405/488/561; voxel size: 1.02 × 1.02 × 2.00 μm^3^; recording interval: 30 min; scale bar: 100 μm.

We observed that in individual organoids the number of cells increases at different rates even if they have similar initial cell numbers. We show that an organoid with an initial cell number of eight increases at a low rate (average: 0.65 cells per hour) and reaches a maximum number of 107 cells after six days, whereas an organoid starting with nine cells increases at a high rate (average: 7.41 cells per hour) and ends up with 1077 cells after six days (**Figure 3a**, **blue and green frames**). Since the splitting procedure results in different sizes of cell clusters, we also analysed one organoid that started with 37 cells and reaches a total number of 644 after six days (average: 4.12 cells per hour) (**Figure 3a**, **red frames**).

To further understand the heterogeneity in proliferation potential and the collective cell behaviour in general, we also quantified the volume and surface area of the organoid as well as the neighbourhood relationships of the single cells (**Figure 3b**). We observed that the organoid with the largest final number of cells did not show the largest volume and surface area (final cell number: 1077, final volume: 10 × 10^7^ voxels, final surface area: 24 × 10^3^ pixels, **Figure 3b, green lines**). In this organoid, the mean number of neighbouring cells was higher within the proximity cell graph (PCG), meaning that cells are neighbours if they are closer than a certain distance (distance: 50 pixels, final PCG-value: 35) in comparison to the organoid with the largest volume and surface area (final PCG-value: 19, final cell number: 644, final volume: 15 × 10^7^ voxels, final surface area: 30 × 10^3^ pixels, **Figure 3b, red lines**). These findings correlate with different cell densities, displayed by the number of neighbouring cells within the Delaunay cell graph (DCG) of 26 and 21 respectively.

Interestingly, we observed frequent size oscillations of single organoids within the culture (**Figure 4c**). Thus, we analysed the size oscillations based on the organoid volume (**Figure 4, definition “size oscillation” see Box 1**). All three organoids showed frequent size oscillations. However, over the time course of six days, the small organoid (**Figure 4a**, **b**, **blue lines**) showed seven oscillations, whereas the two larger organoids showed three (**Figure 4a**, **b**, **red lines**) and two (**Figure 4a**, **b**, **green lines**) events, respectively. We did not observe any correlation between the size oscillation events and the changes in cell number or number and distance of neighbouring cells. Further, we did not observe any size oscillation events in the first 80 hours (within the 5% threshold) nor did we observe any synchronised oscillation behaviour between the organoids within one culture.

**Figure 4:**
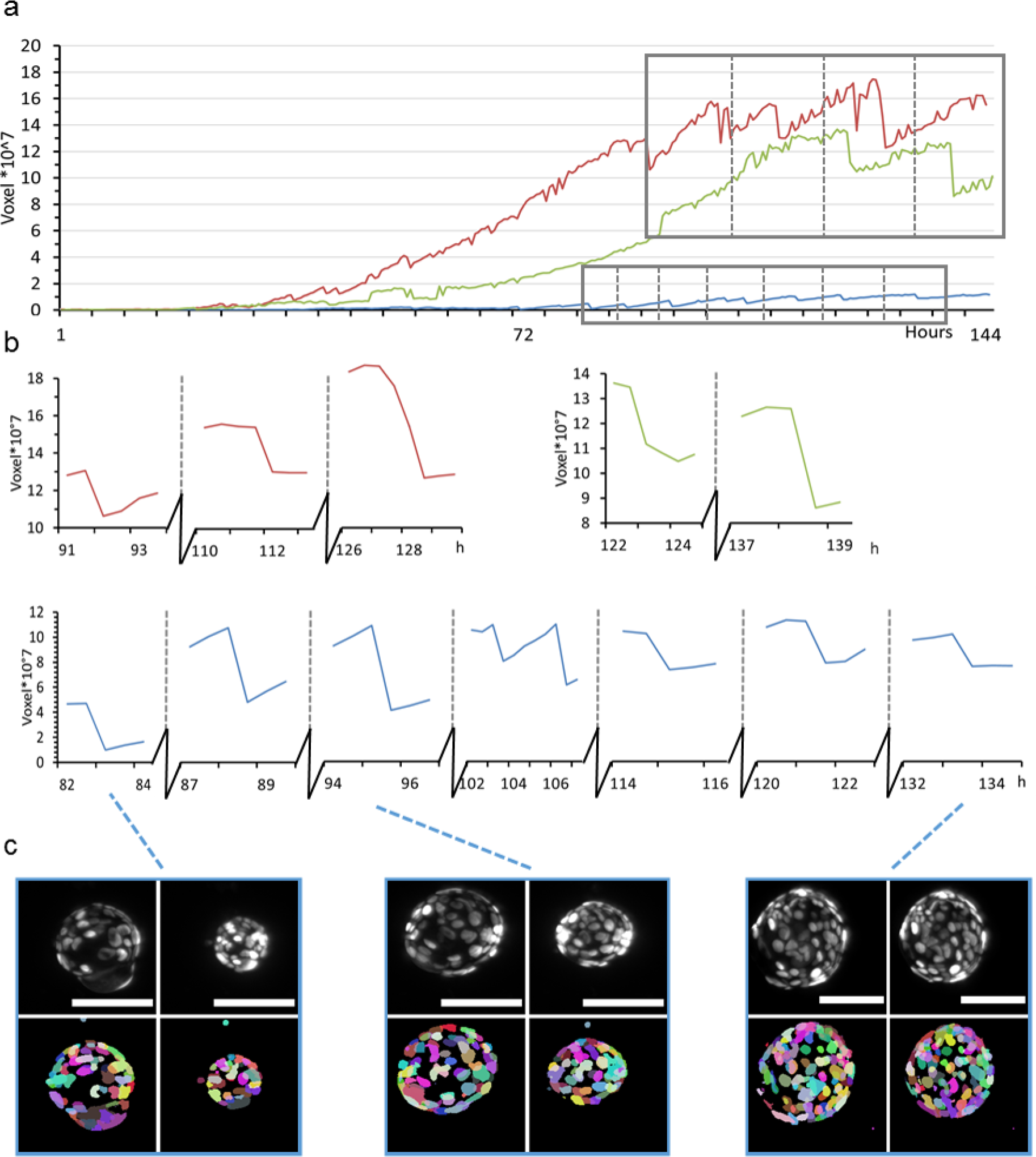
Volume analysis of three representative mPOs reveals different oscillation frequencies. mPOs (seeded and maintained in one Z1-FEP-cuvettes) expressed the nuclei marker Rosa26-nTnG (grey). Organoids were imaged for six days and three representative organoids were analysed with our previously published nuclei segmentation pipeline (Schmitz et al., 2017) in regard to size oscillation events. A size oscillation lasts between 30 minutes and two hours. (**a**) Volume over time for each organoid approximated from the cell nuclei segmentation. (**b**) Detailed analysis of three (red), two (green) and seven (blue) size oscillation events with more than 5% volume reductions. (**c**) Typical images of organoid size oscillation. The upper row shows the nuclei in grey in the raw image, the lower row the segmented cell nuclei. Each colour illustrates a single cell nucleus. Microscope: Zeiss Lightsheet Z.1; objective lenses: detection: W Plan-Apochromat 20x/1.0, illumination: Zeiss LSFM 10x/0.2; laser lines: 561 nm; filters: laser block filter (LBF) 405/488/561; voxel size: 1.02 × 1.02 × 2.00 μm^3^; recording interval: 30 min; scale bar: 100 μm.

### Scaling law derived from simplifying assumptions indicates a dependence of size oscillation events on cell division dynamics

To solve the mechanical principles underlying size oscillation events, we developed a mathematical model based on the following assumptions. Since organoids are spherical single-layer multicellular clusters, they are described by their volume *V*(*t*) and the number of superficial cells *N*(*t*) at time point *t*. We propose a functional relationship for an organoid’s increase in volume 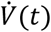, which is derived from two processes: a) The internal pressure of an organoid increases with time, due to an influx following the segregation of an osmotic active substance by the cells. b) Due to mitosis, the cell number *N*(*t*) grows and the surface area *A*(*t*) increases (**Figure 1, Cell division**).

We hypothesise that the increase of the cell number 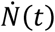 can balance the increase in inner pressure of an organoid and prevent size oscillation events. In the following we show that this requires the cell count *N*(*t*) to grow faster than or equal to *N*(*t*)~ *t*^2^. In return, we expect the occurrence of size oscillations in the case where the cell number increases slower than *N*(*t*)~ *t*^2^. Our estimation is based on the following relations and simplifying assumptions:

i. Organoids form spheres with a volume of

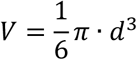

and a surface area of

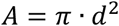 The relation between volume and surface area can be written as

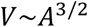
ii. Every cell produces substances, which are secreted into the lumen of the organoid. We assume that the production rate is constant in time and the same for all cells, and therefore proportional to the number of superficial cells.

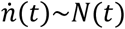
iii. We further assume that the relation between secreted substance and osmotic pressure (Π) follows the van-‘t-Hoff law

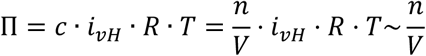
iv. Because of cell division, the surface area grows as a function of time, *A* = *A*(*t*). We neglect cell growth and assume that the cell count *N*(*t*) is proportional to the surface of the organoid, *A*(*t*).
v. The total amount of the substance inside the lumen, *n*, is the accumulated substance produced during organoid growth, therefore

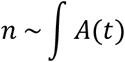

This gives us the relation

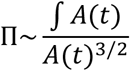

In order to avoid a rupture, the growth of the surface *A*(*t*) has to balance the resulting osmotic pressure Π arising from constant production of *n* by *A*. We can compute the functional form of *A*(*t*) which leads to a constant osmotic pressure Π. We require

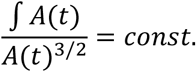

This relationship is fulfilled when *A*(*t*)~*t*^2^. This scaling law provides the following direct implications: A constant cell division rate causes the cell count to increase exponentially. Exponential growth is faster than quadratic, due to the theoretical considerations here we expect no rupture and subsequent size oscillation events. Some of the organoids, however, show a quasi-linear increase in cell numbers, which corresponds to a mitosis rate that is decreasing with 1/*t*. A linear increase is slower than *t*^2^, hence, in these organoids we expect rupture and size oscillations.

In addition, we point out that the surface to volume ratio of a sphere changes with the radius, since the volume grows much faster than the surface. Since the osmotically active substance in the lumen is produced by the surface, this implies that smaller organoids reach a critical internal pressure earlier than large organoids.

### Agent-based mathematical model captures the experimental organoid dynamics and confirms theoretical considerations

In order to confirm our hypotheses we developed a mechanical 3D agent-based model for organoid size oscillations, based on the experimental data obtained by long-term single cell analysis of mPOs (**Figure 5a**; **Supplementary Figure 6**). The model considers intercellular forces (i.e. repulsion and adhesion), internal pressure of the organoids (due to an osmotic imbalance), a bending potential of the cells in order to maintain the spherical shape of the organoid, and cell division. Further, we assume the monolayer of the organoid to break if the mean distance of neighbours exceeds a certain limit.

**Figure 5:**
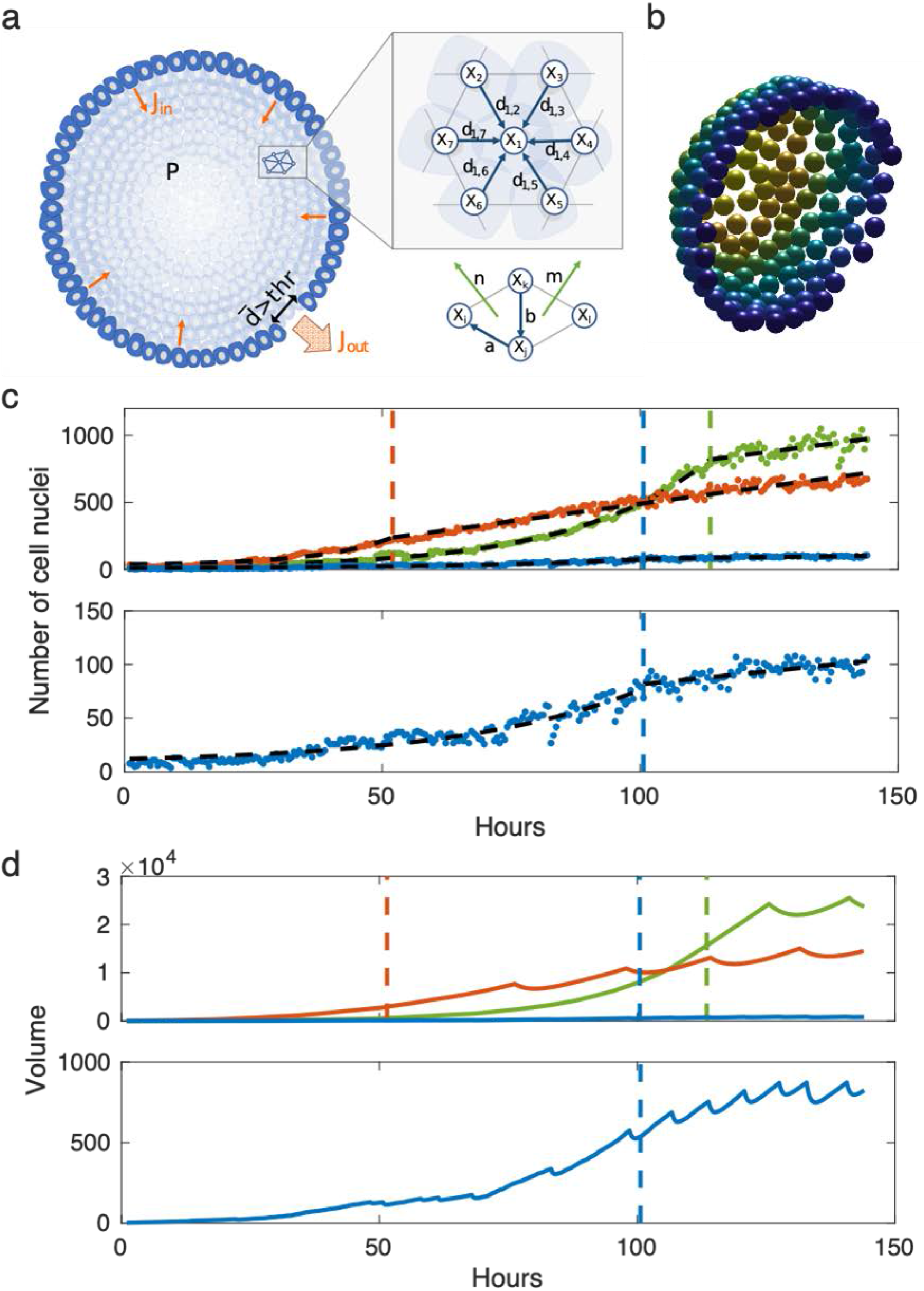
Computational simulation of a multi-agent object. (**a**) Illustration of the general model. (**b**) Snapshot of a simulated sphere cut in half. (**c**) Piecewise exponential-linear fit (black dashed lines) to the growth rates for the long-term single-cell analysis of mPOs. The colours resemble organoids shown in **Figure 3**. The dots indicate the cell count of the organoids. The coloured dashed lines indicate the transition from exponential to linear growth. (**d**) Volumes of the simulated organoids. The coloured dashed lines indicate the transition from an exponential to a linear growth rate. The coloured solid lines show the volumes of the spheres.

We hypothesise that the organoids can be represented as elastic spheres with a growing surface due to cell division (**Figure 5b).**Further, we assume that the cells constantly secrete a substance into the lumen which leads to osmotic influx. This leads to an increase in the internal pressure which, however, can be balanced by an increase in volume. Based on these assumptions, we derived that the organoid can balance the inner pressure when the cell count increases at least quadratically (**Supplementary theoretical considerations**). Furthermore, for small organoids the ratio between surface and volume is smaller than for large organoids. Therefore, small organoids should reach a critical pressure for leakage faster than large organoids. Thus, we expect the size oscillations to critically depend on a) the cell division dynamics and b) the organoid size. The latter (b) is confirmed by the data obtained through bright field analysis (**Figure 6e**).

**Figure 6:**
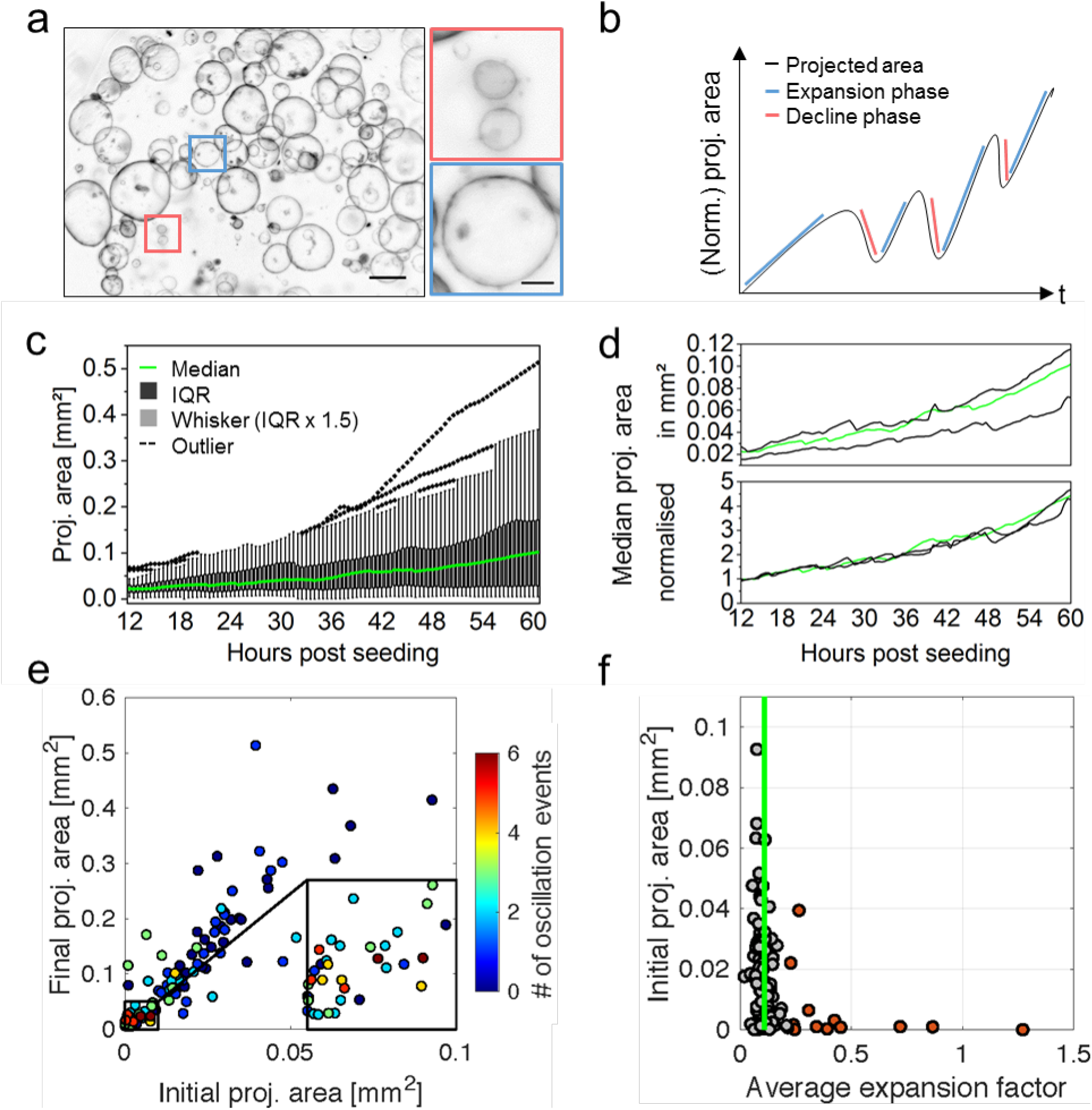
Analysis of multiple monocystic mPOs reveals heterogeneity as well as core regulatory principles. (**a**) Overview bright field images of mPOs displaying a monocystic phenotype. Microscope: Zeiss Axio Observer Z.1; objective lenses: Plan-Apochromat 5x/0.16, avg z-projection, voxel size: 1.29 × 1.29 × 50 μm^3^, scale bar overview: 500 μm, close-up: 25 μm. (**b**) Schematic plot of feature extraction based on time-resolved bright field images. Multiple features, such as the projected luminal area, expansion phases and size oscillation events, were analysed. (**c**) The projected areas of single organoids growing within one well were analysed for 48 hours, starting 12 hours after seeding and revealed high heterogeneity in the projected areas. A high intercultural heterogeneity is illustrated by a broad inter quartile range (black) and outliers (dashed lines) within the box plot (n = 34). (**d**) Medians of projected areas of three wells (technical replicates) differ. The medians of the normalised projected areas are coherent between individual wells. Median shown in (c) is highlighted in green (n = 34, 31, 35). (**e**) The colour code signals the amount of registered size oscillation events. Smaller organoids display an increased number of oscillation events. Close-up reveals location of organoids collapsing four to six times (n = 100). (**f**) The median of the average expansion factor is 0.11 (green line). Independent of their initial projected area [mm^2^], 50% of all organoids display an average expansion factor between 0.09 and 0.14. Here, 12 % of the organoids feature an average expansion factor above 0.22 and are marked as outliers (red) (n = 100).

The model is used to support the theoretical considerations and to qualitatively reproduce the size oscillations of the three analysed mPOs (**Figure 4a-b**). Hereby, the cell division rate is adjusted to match the experimental data. Simulations of two large organoids do not show a size oscillation during phases of exponential cell number increase but start to oscillate after transitioning to a linear growth (**Figure 5c-d**, **green and red lines**). Simulations with a small organoid exhibit size oscillation even during the initial exponential growth, which confirms our hypothesis that small organoids are more prone to rupture and deflation (**Figure 5c-d**, **blue lines**; **Figure 6e**).

Hence, the simulation results show a large qualitatively agreement in the size oscillations with the experimental data and also coincide with the analytical results.

### Time-resolved macro- and mesoscale analysis reveals organoid-to-organoid heterogeneity as well as core regulatory principles

The processes observed using LSFM suggest a vast variety of complex dynamic processes in organoid cultures. In order to analyse the growth characteristics on a macroscale level and to confirm the predictions suggested by the computational model, we established a pipeline based on time-resolved bright field observations. The analysis allows to characterise a culture’s global behaviour. Via semi-automated watershed-based segmentation, the pipeline allows for quantification of the projected luminal areas [mm^2^] over time of several organoids in parallel (**Supplementary Figure 1**). Subsequently, from the normalised projected areas, the relative size increase is evaluated. Further, expansion phases (timing, slope, duration), size oscillation events (timing, slope, duration) as well as minimum and maximum projected luminal areas are identified for individual organoids (**Figure 6b**).

In **Figure 6c**, the projected areas of 34 pancreas organoids growing within one well are plotted over 48 hours. The projected areas illustrate the high heterogeneity, with an area distribution widening over time. After 48 hours of observation, the projected areas have a median of 0.1 mm^2^, while their interquartile range (IQR) ranges from 0.03 to 0.17 mm^2^.

Further, we demonstrate that the bright field pipeline provides consistent and robust growth analysis data in technical replicates (**Box 1**) (three wells, n = 34, 31, 35) (**Figure 6d**). The median values of the normalised projected areas shows no significant differences between the technical replicates (Kruskal-Wallis ANOVA).

The extracted features can further be used in downstream analyses to categorise organoid behaviour. We found that the size of an organoid is crucial for the number of oscillation events it displays. Consistent with the LSFM data and the mathematical model, the bright field analysis shows that initially smaller organoids (area < 0.01 mm^2^) feature more size oscillation events, while initially larger organoids display less oscillation events (area > 0.01 mm^2^) (**Figure 6e**). Besides that, the average expansion factor (a measure for expansion speed consistency) with a median average value of 0.11 is similar between organoids with various initial areas −50% of all values range between 0.09 and 0.14, while only 12% of the evaluated organoids are outliers with values above 0.22 (**Figure 6f**). Further, the linear correlation between the initial area and the final area becomes apparent (R^2^ = 0.7445), which shows that the growth is independent of the initial area (**Supplementary Figure 7a**). This indicates strong similarities in expansion speed consistency between individual organoids within one culture despite their (high) size heterogeneity.

Besides the already mentioned features, other extracted features facilitate the definition of quantitative reference parameters of organoid systems. By comparing the final area to the maximum area, for example, continuous growth of mPOs during the analysed time window is proven. A comparison of the initial area to the minimum area identifies size oscillation events or overall descending size progression within organoid cultures. In mPOs, the minimum area falls only slightly below the initial area, which can be associated with oscillation events (**Supplementary Figure 7c**).

Besides the average expansion factor, analysis of the maximum expansion factor indicates expansion speed variations within organoid cultures. As a variable factor, the maximum expansion can be used to compare different culture conditions (**Supplementary Figure 7b**). An additional feature, which is likely to change upon differentiation or other perturbations (e.g. drug treatment), is the organoid circularity. In healthy mPOs, the circularity is 0.9 on average and the deviation around the average narrows over time (**Supplementary Figure 7e)**. In addition to the analysis of monocystic epithelial organoids like mPOs, our bright field pipeline can also be used to analyse deviating organoid morphologies like polycystic hCCOs (**Supplementary Figure 8a-f**). Polycystic hCCOs show an average circularity of 0.8 over the course of 48 hours of observation (**Supplementary Figure 8b**). Therefore, as a general culture feature, the circularity can serve as an additional quality control parameter.

## Discussion

We describe two complementary light sheet and bright field imaging pipelines for the time-resolved, multiscale quantitative analysis of single-cell and collective cell behaviours in organoids. The goal is the characterisation of the heterogeneity of organoid cultures in their entirety.

The light sheet pipeline led to the identification of several dynamic processes typical of organoid cultures on micro- (single-cell) and mesoscale (individual organoid) levels. The resolution and contrast of the light sheet images allowed the quantification of these processes with nuclei segmentation and single-cell tracking. The bright field pipeline allowed quantifying the dynamics of individual organoids as well as of entire organoid cultures on a macroscale level. In both pipelines, we choose a time resolution of 30 minutes to capture the growth behaviour of organoids. We selected this time interval as a technical compromise in the need for a high temporal resolution to resolve dynamic processes (e.g. budding or cell rearrangement processes (Serra et al., 2019)), and the need to observe these processes over long observation periods (e.g. to study cell differentiation (Fatehullah et al., 2016; Lancaster et al., 2013).

In previous studies, chromosomal segregation errors in organoids were monitored using confocal or spinning disc microscopy, capturing single organoids in a z-range of 60 μm (3-4 min intervals) (Bolhaqueiro et al., 2019). The observation of single mitotic events with LSFM over multiple days, from the initial seeding to the plateau-phase of growth, can therefore increase the throughput and allow the analysis of larger organoids to monitor single cell behaviour. Serra *et al.* used an inverted LSFM to analyse the development of single organoids originating from single cells. By parallelisation, they were able to image multiple organoids (Serra et al., 2019). Our light sheet pipeline combines a parallelised acquisition of more than 100 organoids, within a volume of up to 8 mm^3^ given by the Z1-FEP-cuvette, with a high spatial (1000 z-planes, 2 μm spacing), as well as a high temporal resolution and still allows long-term observations.

Our analysis of mPOs and hCCOs reveals their highly heterogeneous and multi-faceted growth patterns and common morphological dynamics independent of their carcinogenic or healthy origin. This matches the observation of intrinsic abilities of single intestinal organoid cells to form asymmetric structures (Serra et al., 2019), as well as former studies that have not addressed heterogeneity directly, but already showed variable organoid sizes and irregularly occurring rupture events (Mahe et al., 2013; Schlaermann et al., 2016; Schwank et al., 2013; Sebrell et al., 2018).

Adult tissue-derived organoids can develop from single cells or cell clusters, although starting from single cells results in lower organoids formation efficiency, which hampers the systematic analysis of cellular behaviours (Serra et al., 2019). Starting from cell clusters, the cell survival and therefore the multiplication-rate of cell material is higher. The production of large amounts of material for a potential clinical application is therefore ensured (Dossena et al., 2020). However, starting from clusters, the heterogeneity of the cultures dynamics increases. Since our pipelines capture both aspects, they support the understanding of clonal formation as well as the determination of quality control parameters for clinical applications of organoids. We illustrate at multiple scales that frequent size oscillations in mPOs are common during the period of growth. It occurs more often in smaller organoids, showing that the initial size of the cell cluster is a crucial factor. This observation is in contrast to previous findings by Sebrell *et al.* who observed a trend to more size oscillation in large (> 200 μm) organoids. (Sebrell et al., 2018). Since they analysed human gastric epithelial organoids, this raises the question of how organoid systems differ in their behaviour and growth even if they are both of epithelial origin and grown under similar culture conditions. This further illustrates the need to define core regulatory principles of organoid systems.

Furthermore, we observed and defined the fusion process of organoids. This process has never been described for organoids in 3D in comparable temporal and spatial resolution before and shows similarities to several processes in mammalian embryonic development (Kim et al., 2015) and organ maintenance (Bruens et al., 2017). Kim *et al.* analysed the fusion of the palatal shelfs during craniofacial development and identified three general stages for tissue fusion, which are comparable to our observed organoid fusions. It is initialised by the convergence of two epithelial layers, migratory movements towards one epithelial layer and subsequent rupture of the single cell layer. Dumortier *et al.* showed that the formation of the blastocoel during mouse pre-implantation is a result of a highly dynamic process of cell-cell ruptures and fusion events that lead to a final large lumen (Dumortier et al., 2019). They predict that hydraulic fracturing of cell-cell contacts guides the rupturing process, which is consistent with our mechanical 3D agent-based model and our data. Observing size oscillation or fusion events of organoids in high temporal and spatial resolution provides an *ex vivo* model to understand dynamic processes like tissue fusion or cavity development.

The multi-faceted dynamic behaviour of organoids is reflected in their motion. We were able to show that organoids differ in their overall rotation speed and rotation direction, in their cell motion depending on their position within the organoid and that not all organoids show rotational behaviour. In general, rotation in organoids is poorly described directly, however, dynamic processes are already investigated in other 3D cell culture systems (Ferrari et al., 2008; Hirata et al., 2018; Marmaras et al., 2010; Tanner et al., 2012). Sebrell *et al.* related the rotation of gastric epithelial organoids to the passage number and patient/donor history (Sebrell et al., 2018). Wang *et al.* investigated the rotation of 3D human mammary epithelial acini and identified that cell polarity and microtubules are essential for rotation (Wang et al., 2013). It remains an exciting question, why not all investigated organoids show rotation.

Computational and mathematical *in silico* models are a valuable tool to understand the underlying mechanics of 3D cell culture behaviour (Eils et al., 2013). They can be used to predict organoid behaviour in conditions that are challenging to implement in experiments or when perturbations of normal conditions occur (Dahl-Jensen et al., 2017; Eils et al., 2013). However, only relatively few models for organoid systems have been developed (Montes-Olivas et al., 2019).

Here, we implemented a mechanical 3D agent-based model that relies on a limited set of assumptions (namely intercellular forces, internal pressure of the organoid, bending energy of the surface and cell division). We showed that the model is a valuable instrument for the description of spatiotemporal dynamics of organoids. We were able to recreate the qualitative growth curve of the three segmented organoids and showed that the frequent size oscillation of organoids is not directly associated with mitosis (for further experimental analysis of the mechanisms underlying organoid size changes see also Yang *et al.* (Yang et al., 2020). Instead, the model indicates that the decline process relies on cells losing cell contacts due to mechanical stress exerted by the internal luminal pressure (Ruiz-Herrero et al., 2017). Further, the model confirms that the size oscillation dynamics are dependent on the organoid volume-to-surface ratio and its dynamics with exponential and linear growth phases. The disagreement between the simulation and data concerning the volume of the medium and large organoid results from the fact that the cell size differs between the two organoids. The large organoid exhibits a higher cell density, which implies a smaller cell size compared to the medium size organoid. Different cell sizes are not considered in the current version of the model but can be included to reflect these different phenotypes.

Our light sheet pipeline shows that the small organoid has a higher size oscillation frequency (**Box 1**) than the larger organoids. The theoretical considerations and the mathematical model support this observation: size oscillations are affected by an increased surface-to-volume ratio. The bright field pipeline further confirm this observation. In summary, the simulation of epithelial organoid growth predicts organoid behaviour and helps to understand the intrinsic mechanisms responsible for the organoid phenotype. Further, it is straightforward to generate a cost- and time-effective tool to predict possible outcomes of external stimuli like drug treatments for instance (Montes-Olivas et al., 2019).

The bright field pipeline enables the quantification of culture dynamics on meso- and macroscale level, generating robust data on organoid growth behaviour and allowing the quantification of heterogeneity in whole organoid cultures. The pipeline has been extensively applied in the LSFM4LIFE (www.lsfm4life.eu) and the Onconoid Hub projects to measure and optimise the growth of human pancreas-derived organoids in synthetic hydrogels and identify novel drug candidates for the treatment of intrahepatic cholangiocarcinoma (manuscripts in preparation). In perspective, the same analysis is suitable to determine parameters of organoid growth for stem cell therapy (Aberle et al., 2018; Huch et al., 2017; Lancaster et al., 2019; Nagle et al., 2018) and to characterise patient-specific responses for optimising personalised drug treatments or assaying the onset of resistance in cancer therapy (Broutier et al., 2017; Fan et al., 2019; Nagle et al., 2018; Ooft et al., 2019).

In conclusion, our multiscale analyses of diverse organoid cultures have a great potential for further investigations of epithelial organoids and many other complex culture systems.

## Material and methods

### Organoid culture

hCCOs were initiated from primary liver tumour biopsies of cholangiocarcinoma patients (~0.5 cm^3^) collected during surgery performed at the Erasmus MC – University Medical Center Rotterdam (NL) and cultured as previously described (Broutier *et al.*, 2017). mPOs were obtained from Meritxell Huch (Gurdon Institute, Cambridge, UK) and cultured as described (Huch et al., 2013).

#### Transgenic murine pancreas-derived organoids

The transgenic mPO line was obtained from Meritxell Huch’s laboratory at the Wellcome Trust/CRUK Gurdon Institute, University of Cambridge, UK. The cells were isolated from the adult pancreas of the Rosa26-nTnG mouse line (*B6;129S6-Gt(ROSA)26Sortm1(CAG-tdTomato*,-EGFP*)Ees/J*, stock no. 023035, The Jackson Laboratory, Bar Harbor, Maine) according to the isolation protocol (Broutier et al., 2016).

#### Viral transduction of human liver-derived tumour organoids

hCCOs were transduced using a third generation lentivector (pLV-Puro-EF1A-H2B/EGFP:T2A:LifeAct/mCherry, vector ID: IK-VB180119-1097haw, custom-made by and commercially obtained from AMSBIO, Abingdon, UK) for stable expression of the fluorescent fusion proteins H2B-eGFP to visualise cell nuclei and LifeAct-mCherry to visualise the F-actin cytoskeleton. Lentiviral particles were commercially obtained from AMSBIO (Abingdon, UK). Viral transduction of organoids for stable expression of fluorescent markers in hCCOs was performed according to a protocol published by Broutier *et al*. (Broutier *et al.*, 2016) with slight modifications. In brief, organoids were dissociated into small cell clusters by mechanical fragmentation in pre-warmed (37°C) trypsin and subsequent incubation for 5-10 min. All centrifugation steps were carried out at room temperature. Positive (transduced) organoids were selectively picked under semi-sterile conditions instead of being selected by puromycin administration, and were expanded into positively labelled cultures without sorting.

#### Ethical approvals

The Ethics Approval REC No. 12/EE/0253 from the UK National Research Ethics Service (NRES) covers the ethical issues involved the generation and culture of the murine pancreas-derived organoids used in the LSFM4LIFE project and in this work. Medical ethical approval for the use of patient liver tumour biopsies for research purposes has been granted by the Medical Ethical Committee (METC) of the Erasmus Medical Center in Rotterdam, The Netherlands (MEC-2014-060). Patients provided written informed consent and all methods were performed in accordance with the relevant guidelines and regulations.

### Light sheet pipeline

In order to generate single-cell resolved high-content data of organoid dynamics, the previously published ultra-thin FEP-foil cuvette (Hötte et al., 2019) was used. To implement it into the Zeiss Lightsheet Z.1 system (Carl Zeiss AG, Oberkochen, Germany) a new positive module was produced, and the cuvette was connected with a capillary.

#### Fabrication of positive moulds for vacuum forming

We designed positive moulds of the cuvettes for the use in the Zeiss Lightsheet Z.1 system by using the free CAD software “123D Design” (version 2.2.14, Autodesk). We 3D-printed the positive moulds by using the service of the company Shapeways. Before use, the positive moulds were inspected by stereomicroscopy and cleaned by immersion in an ultrasonic bath.

#### Cuvette fabrication with vacuum forming

For a detailed description of the cuvette fabrication with vacuum forming refer to Hötte *et al.* 2019 (Hötte et al., 2019). In brief, a 10 cm x 10 cm squared patch of FEP-foil (50 μm thickness, batch no. GRN069662, Lohmann Technologies, Milton Keynes, UK) was clamped into the vacuum-forming machine (JT-18, Jin Tai Machining Company), heated up to 280°C and pressed onto the square cross-section positive mould described in **Supplementary Figure 2d-e**. The positive mould was assembled with a 2 mm x 2 mm 3D-printed square cross-section rod and a glass capillary (borosilicate glass 3.3, material no. 0500, Hilgenberg GmbH, Malsfeld, Germany), cut to a length of about 15 mm with a diamond cutter (**Supplementary Figure 2d-e**). After vacuum forming of the FEP foil, the square cross-section rod was carefully removed, leaving the FEP cuvette supported by the glass capillary, which serves as mechanically stable connection with the Zeiss Z.1 holder. A shrinking tube (flame retardant polyolefin tube, size 3, cat. no. E255532, G-APEX, Yuanlin City, Taiwan) was used to close the FEP cuvette and to connect it to the glass capillary connected with the Zeiss Lightsheet Z.1 xyz stage (Blaubrand intraMark 200 μl micropipette, cat. no. 708757, BRAND GmbH & Co. KG, Wertheim am Maim, Germany) (**Supplementary Figure 2f**). In order to pipette the samples into the cuvette, the capillary was removed. Finally, the complete FEP cuvette setup was cleaned with a detergent solution (1% Hellmanex-II in ultrapure water), sterilised in 75% Ethanol for at least two hours and washed twice with PBS.

#### Specimen preparation

Organoids were cultured as described. During the splitting procedure 20 μl of ECM containing mPO/hCCO cell clusters were filled into the cuvette. To avoid air bubbles the use of a 20 μl pipette tip is recommended (TipOne 10/20 μl, STARLAB, Hamburg, Germany). Subsequently, a glass capillary (Blaubrand intraMark 200 μl micropipette) that has been filled with expansion medium beforehand was connected. To ensure no leakage, the connections between the shrinking tube, the FEP cuvette and the glass capillary were wrapped with Parafilm (**Supplementary Figure 2**).

For imaging, the FEP cuvette attached to the capillary, was inserted in the Zeiss Lightsheet Z1. The imaging medium in the Zeiss Lightsheet Z1 chamber was DMEM (without phenol red) dosed with 2% penicillin and streptomycin and HEPES (1:100). During the time of observation, the medium exchange was conducted under semi-sterile conditions directly at the microscope with a 10 μl microloader tip (Microloader Tip 0,5-10 μl / 2-20 μl, Eppendorf AG, Hamburg, Germany) every two days.

#### Image acquisition and microscopic feature extraction

Image stacks of the entire Z1-FEP-cuvette containing the mPOs/hCCOs were acquired with the Zeiss Lightsheet Z1 microscope. The mPO cells expressed Rosa26-nTnG and were exited with a 561 nm laser. The hCCO cells expressed H2B-eGFP and LifeAct-mCherry and were exited with a 488 nm and 561 nm laser. Both cell lines were imaged with a Carl Zeiss W Plan-Apochromat 20x/1.0 UV_VIS objective and illuminated from two sides with Zeiss LSFM 10x/0.2. During the image acquisition, the chamber was temperature and CO_2_ controlled and constantly filled with pre-warmed DMEM containing 2% penicillin and streptomycin and HEPES (1:100). mPO image stacks were cropped towards the corresponding organoid (Fiji, ImageJ) and all single time points of each organoid were segmented and processed separately by using the previous published multiscale image analysis pipeline (Schmitz et al., 2017) with the configurations mentioned in **Supplementary Table 1**. For feature extraction and the surface approximation, the configurations mentioned in **Supplementary Table 1**were used.

#### Arivis Vision4D

3D volume rendering and 3D cell tracking was performed with Arivis Vision4D (Version: 3.1.3, Arivis AG, Munich, Germany). Prior to segmentation, image stacks were filtered with “Particel enhancement” (Diameter: 10, Lambda: 1). Single cell nuclei were subsequently segmented with “Blob Finder” (Segment value: 500 μm, Threshold: 5, Watershed level: 1.303, NormalizePerFrame: true, SplitSensitivity: 82%) and tracked with “Segment Tracker” (Motion type: Brownian Motion (centroid), Max. distance: 5 μm, Track: Fusion: false – Divisoins: true, Weighting: Multiple).

#### Fiji/ImageJ

Organoid and nuclei sizes for the visual inspections part of the results were measured manually on maximum intensity z-projections of the acquired fluorescence image data using FIJI/ImageJ.

### Mechanical 3D agent-based model

An individual cell-based model was implemented to explain the size oscillations of the pancreas-derived organoids. The mathematical model was given as a set of stochastic differential equations that were solved using the Euler-Maruyama method.

To describe the pancreas-derived organoid, we assumed it has a roughly spherical shape, with cells forming a monolayer filled with fluid at a different pressure relative to the environment. The volume of the organoid is affected by two mechanisms: a) the influx of liquid caused by an osmotic imbalance or active pumping of the cells, and b) cell division. While a) is increasing the internal pressure, b) leads to a relaxation of the surface.

Each cell was described by a small set of features, i.e. a position in 3D space and a cell size denoted by its radius. Displacement of the cells was described as a response to three forces: 1) external forces exerted by surrounding cells, given as a spring potential, 2) internal pressure of the organoid pushing the cells outwards, given by the ideal gas law, and 3) a surface bending energy, keeping the organoid in its spherical shape.

Cell division was adjusted to match the experimental data, obtained by long-term single-cell analysis of pancreas-derived organoids, but can easily be adapted to other growth dynamics. If the average distance of neighbouring cells exceeds a certain limit, we assumed the mechanical stress to be too high and a leakage in the shell of the organoid emerges. Through the rupture, the internal liquid is released and the internal pressure decreases. Thus, the mechanical forces, exerted on the cells might relax and the organoid deflates. When the average distances between all neighbouring cells falls below a given threshold, the shell closes and the liquid stops to be released.

For a more detailed description of the model, we refer the reader to the supplementary material.

### Bright field pipeline

#### Specimen preparation

For the bright field analysis, organoids were seeded in 25 μl ECM (Matrigel, Corning, New York) droplets in suspension culture plates (48-well, Greiner Bio-One, Kremsmünster, Austria), overlaid with 250 μl expansion medium and cultured for 12 h before imaging. They were then imaged every 30 min in a 3×3 tile imaging (15% overlap) mode using the Zeiss Cell Observer Z.1, fully equipped with an incubation chamber and motorised stage using a Plan-Apochromat 5x/0.16 objective, with a pixel size of 1.29 μm x 1.29 μm. In total, ten planes throughout the droplet were imaged, with a z-distance of 50 μm (mPOs) and 65 μm (hCCOs), respectively, capturing a z-range of 450 to 585 μm.

#### Image processing and organoid segmentation

Organoid growth rates were determined using a python custom-made pipeline for bright field-based image segmentation. The whole pipeline was equipped with a general user interface. The recorded time-lapse image stacks were pre-processed with Fiji (ImageJ version 1.51n, Java version 1.8.0_6 (64-bit)) by reducing the dimensionality of the raw data set from 9 (3×3) tiles with 10 z-planes each to one stitched image with one z-plane per time frame using the functions Average Intensity ZProjection, Subtract Background (Rolling ball radius: 700 pixels, Light background, Sliding paraboloid, Disable smoothing) and Grid/Collection stitching (Preibisch et al., 2009) (Type: Grid: row-by-row, Order: Right & Down). The resulting image stacks were further subjected to filtering (Median, Radius: 5 pixels), background subtraction (Rolling ball radius: 500 pixels, Light background, Sliding paraboloid), and the projected luminal areas of the organoids were using the Fiji plugin Morphological Segmentation (MorphoLibJ (Legland et al., 2016) → Segmentation → Morphological Segmentation; Border Image, Tolerance: 10 (mPOs), 12 (hCCOs), Calculate dams: true, Connectivity: 6). Segmented luminal areas were measured with the Fiji plugin Region Morphometry (MorphoLibJ (Legland et al., 2016) → Analyze → Region Morphometry).

The results were plotted and statistically evaluated (Kruskal-Wallis ANOVA, p < 0.05) using OriginPro 2019 or Excel. For a normalisation, the projected areas were normalised to the median of the fifth time point.

#### Mesoscopic feature extraction

Quantitative features were extracted using a Python script and were defined as follows: A size oscillation event consists of a decline phase followed by an expansion phase. The start of a decline phase was defined as the time point after which the area declines by 5%, and the end is marked if the area increases again. Expansion phases were defined between the end of a decline phase and the start of the following decline phase. As additional criterion, the duration of expansion phases is greater than or equal to five time points, and the correlation coefficient of the fitted polynomial is above 0.9. The number of decline and expansion phases per organoid was determined including their duration and slope. Subsequently, maximum and average expansion slopes were computed. The average expansion factor is specified as the average slope of all detected expansion phases per organoid. The maximum expansion factor is specified as the maximum slope of all detected expansion phases per organoid. Outliers in average expansion were defined as smaller than the first quartile minus 1.5 x IQR or above the third quartile plus 1.5 x IQR. The circularity was monitored continuously and is defined as 4π(area/perimeter^2^). Its standard deviation is displayed as the average standard deviation in all analysed wells. Organoids displaying a circularity below 0.6 were considered as deficiently segmented and were excluded from further analyses. Due to deficient segmentation during organoid formation the projected area was normalised to the fifth time point of acquisition.

##### Box 1.

###### Box 1.

• **Agent-based model:** A computational model, in which cells are represented as autonomous decisionmaking agents. Agents interact with their environment based on a given ruleset. Agent-based models allow fora high level of physical detail and can reproduce complex behaviour patterns.
• **Biological replicate:** An independently performed experiment with either a new organoid line ora complementary passage ofthe same organoid line.
• **Core regulatory principles:** Mechanistic understanding of multicellular systems (Sasaiefa/., 2013).
• **hCCO:** Human cholangiocarcinoma-derived organoids.
• **mPO:** Murine pancreas-derived organoids.
• **Monocystic:** Spherical mono-layered epithelium with a single lumen.
• **Multiscale organoid analysis:** The analysis of an entire organoid culture at the three levels of observation:

- **Microscale:** Single cell
- **Mesoscale:** Individual organoid
- **Macroscale:** Entire organoid culture
• **Non-invasive imaging:** Organoids are imaged without the use of any additional fluorescent dye, any additional substance or any chemical-physical influence besides the medium exchange to ensure optimal growth conditions.
• **Organoid degeneration:** Overall shrinking ofthe organoid accompanied by nuclear condensation and fading nuclei signals.
• **Organoid luminal dynamics:** Formation, retraction and rupture of duct-like structures within the lumen of an organoid.
• **Organoid size:** The term “size” refers to an organoid’s volume [voxels] based on surface approximations derived from the light sheet image data or to the organoid’s projected luminal area [mm^2^] based on segmented equatorial planes derived from the bright field image data.
• **Polycystic:** An irregularly shaped multi-layered epithelium surrounding multiple lumens.
• **Projected luminal area:** The segmented and measured equatorial plane of an organoid.
• **Size-oscillation event:** Size alteration of an epithelial organoid characterised by the following phases:

- **Decline phase:** Starts with the time point, upon which the volume/area is 5% smaller than at the previous time point. The decline phase ends when volume/area increases again.
- **Expansion phase:** Starts with the time point at the end of a decline phase and ends the start ofthe following decline phase if this phase comprises more than five time points.
• **Size-oscillation frequency:** The rate with which size oscillation events occur.
• **Technical replicate:** One well of organoids that contains several individual organoids of identical origin.
• **Z1-FEP-cuvette:** Custom sample holder for live imaging with the Zeiss Lightsheet Z.1 microscope system based on the previously described ultra-thin FEP-foil cuvette (Hötte *et al.*, 2019).

## Acknowledgements

FP, EHKS, LH, TM, MKoch thank the EU Horizon2020 project LSFM4LIFE (grant no. 668350-2), the ZonMw-BMBF joint sponsored project Onconoid Hub (grant no. 114027003), and the DFG Cluster of Excellence Frankfurt “Macromolecular Complexes” (CEF-MCII) for funding.

FM is generously supported by the Giersch foundation. TL, FM are supported by funding from the Hessen State Ministry for Higher Education, Research and the Arts in the framework of the Loewe Program (DynaMem, CMMS).

## Authors’ contribution

LH and TM cultured the murine pancreas-derived organoids. LH imaged, analysed and evaluated the data acquired with the bright field pipeline. TM imaged, analysed, 3D rendered and evaluated the data acquired with the light sheet pipeline. TM evaluated the pH and temperature properties of the Zeiss Lightsheet Z.1 microscope system. MKoch designed the viral transduction vector and cultured the human liver-derived tumour organoids. TL, MKurtz and FM developed the 3D agent-based mathematical model. TL analysed and visualised the data from the experiments and simulations. MKurtz and FM derived the mathematical relations on the dependence between pressure and cell division dynamics. JT developed the bright field pipeline and generated a general user interface. TL commented and improved the bright field analysis pipeline. SML transduced the human liver-derived tumour organoids. MMAV and LJWvdL provided the human liver-derived tumour organoid cultures. MH provided the murine pancreas-derived organoid cultures. FP invented the ultra-thin FEP-foil cuvettes, designed and improved their fabrication process. FP and TM adapted the ultra-thin FEP-foil cuvettes for the application to the Zeiss Lightsheet Z.1 microscope. FM, FP and EHKS supervised the work. LH, TM, MKoch, TL, MKurtz, FM and FP wrote the manuscript. All authors contributed to the writing process, and revised and approved the manuscript.

## Supplementary Information

### Supplementary Figures

**Supplementary Figure 1:**
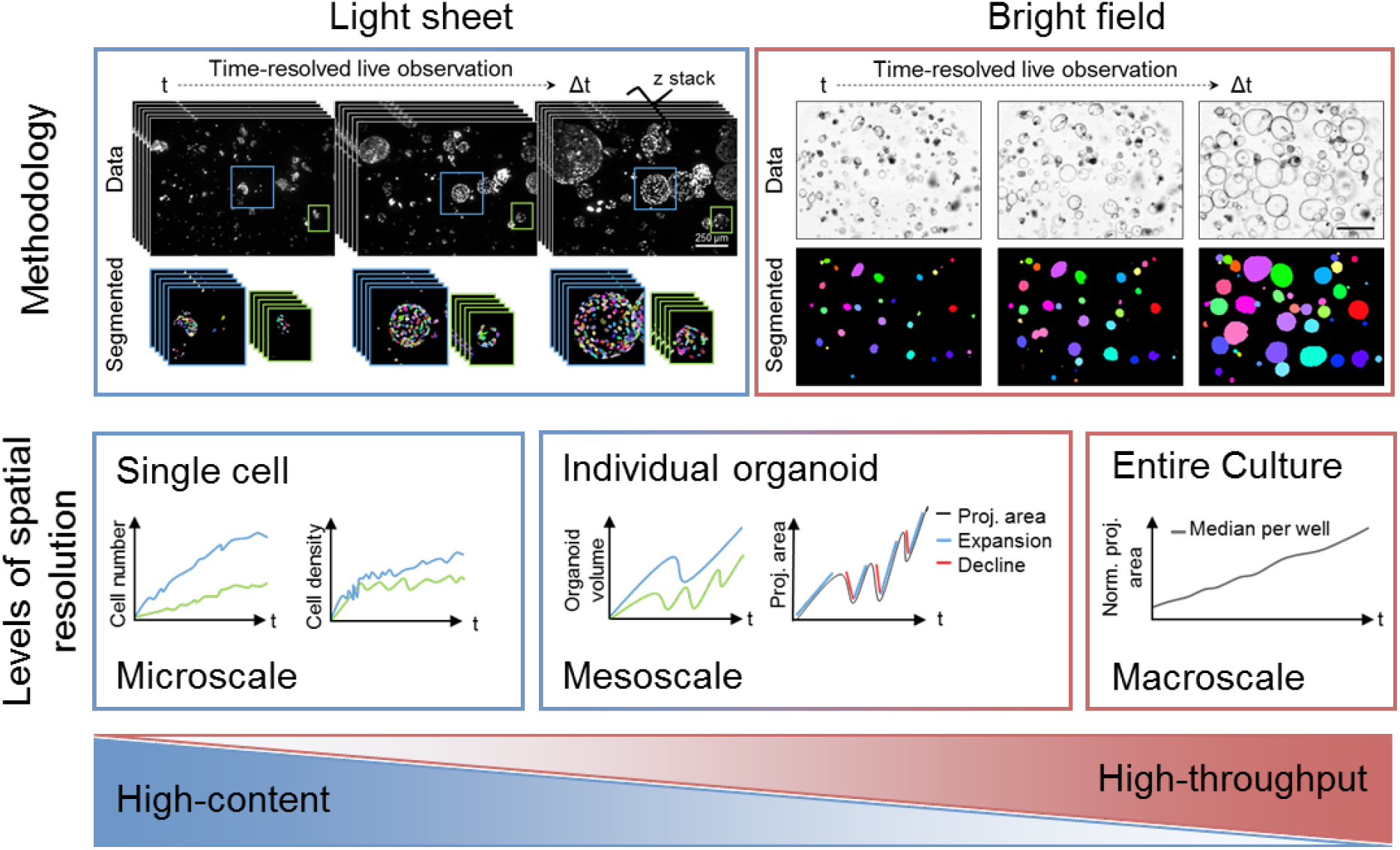
Light sheet and bright field time-resolved observation allow quantitative analyses of micro-, meso- and macroscale dynamics in organoid cultures. Using a light sheet-based fluorescent microscope time-resolved image stack of organoids are recorded. The high-resolution images are subjected to nuclei segmentation (Schmitz et al., 2017) for the quantification of dynamics on single cell level (microscale). Besides that, dynamics, such as size oscillation events, of individual organoids (mesoscale) can be analysed. The restricted throughput of this pipeline is matched with the analyses based on time-resolved bright field images. Here, the dynamics of high numbers of organoids are quantified based on the normalised (norm.) projected (proj.) luminal areas. The pipeline also enables the observation of entire organoid cultures (macroscale) within individual wells.

**Supplementary Figure 2:**
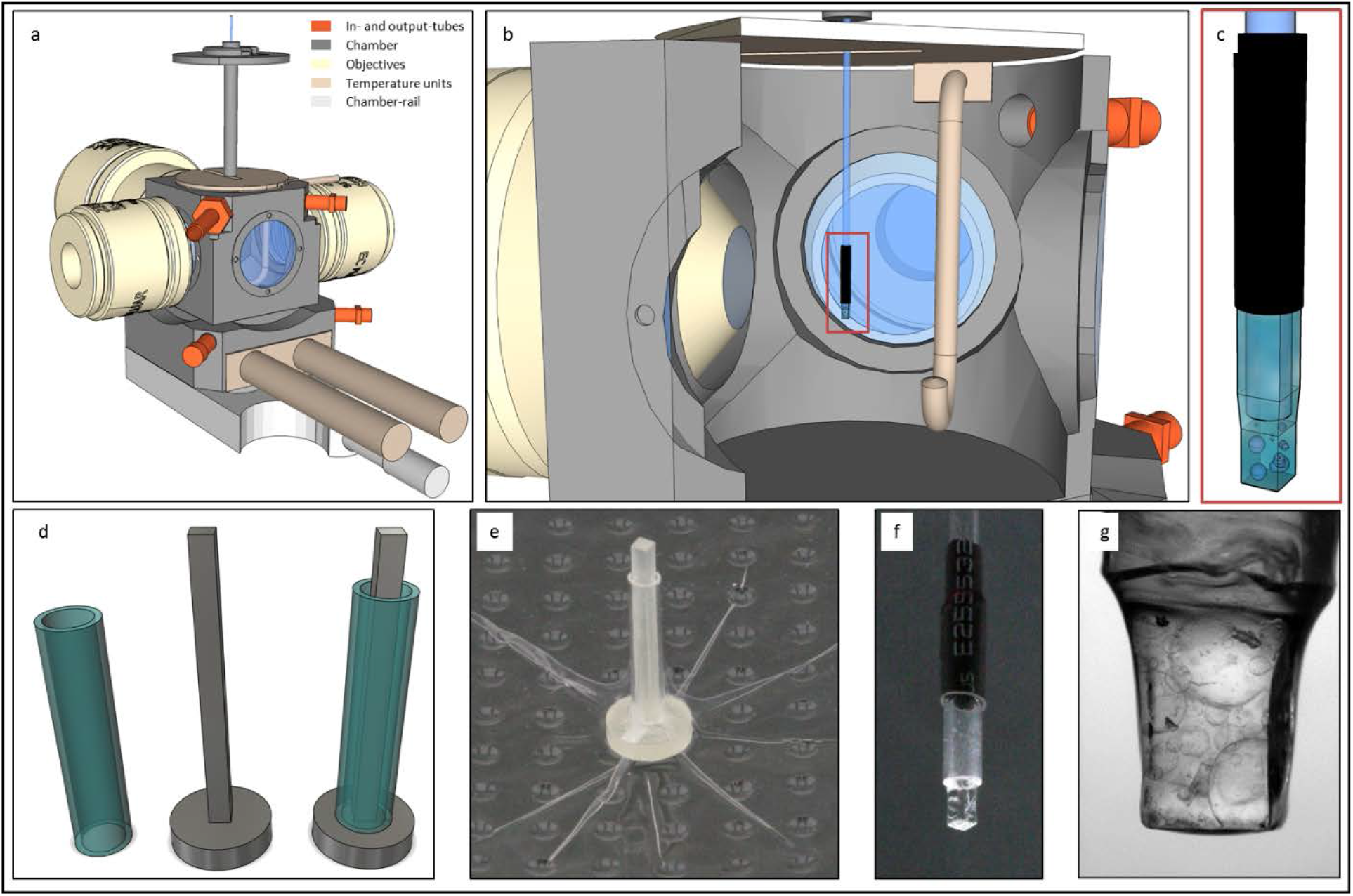
Ultra-thin FEP-foil cuvette holders for live recordings with the Zeiss Lightsheet Z.1 microscope system - Z1-FEP-cuvettes. (**a**) Illustration of the general setup of the Zeiss Lightsheet Z.1 microscope. (**b**) Close-up of the microscope chamber with the downwards directed Z1-FEP-cuvette enclosing the sample. (**c**) Close-up of the sample holder. The shrinking tube that seals the FEP cuvette and connects it with the glass capillary is depicted in black. (**d**) CAD-derived drawings of positive moulds of the FEP cuvette and the glass capillary needed to produce the Z1-FEP-cuvette. (**e**) Printed mould with a glass capillary used to form the Z1-FEP-cuvette in the vacuum forming process. (**f**) Ready-to-use Z1-FEP-cuvette. (**g**) mPOs grown for 7 days in the Z1-FEP-cuvette.

**Supplementary Figure 3:**
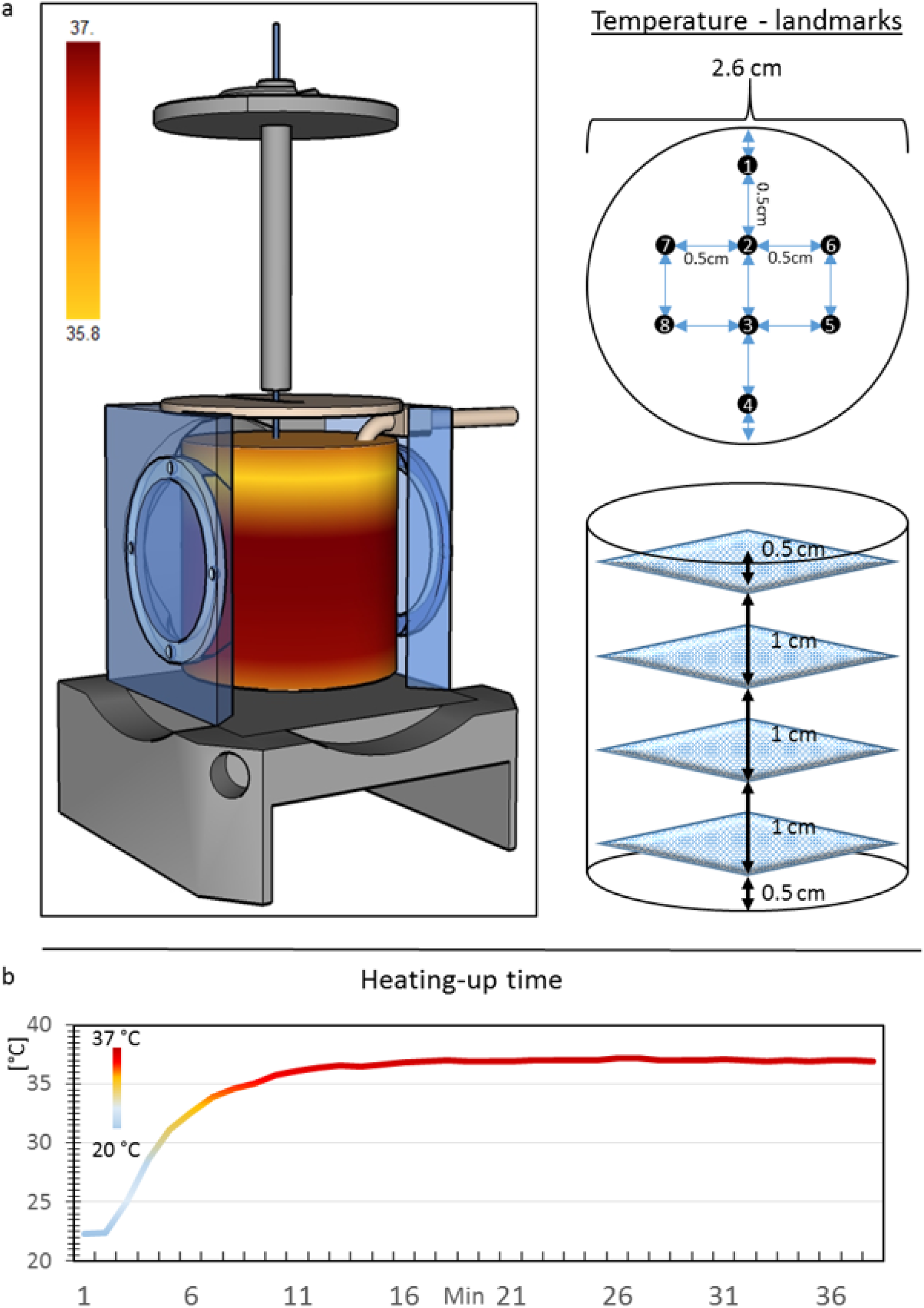
Validation of the temperature properties of the Zeiss Lightsheet Z.1 microscope. (**a**) Illustration of the temperature distribution inside of the Zeiss Lightsheet Z.1 microscope chamber and the corresponding measurement landmarks. Beside the open, upper part with a slightly lower value, the temperature is equally distributed throughout the chamber. (**b**) Results of the measurement of the heating-up time. The included heating unit of the microscope needs to heat up the medium starting from room temperature (21°C). After 12 minutes the medium reaches the physiological temperature of 37°C..

**Supplementary Figure 4:**
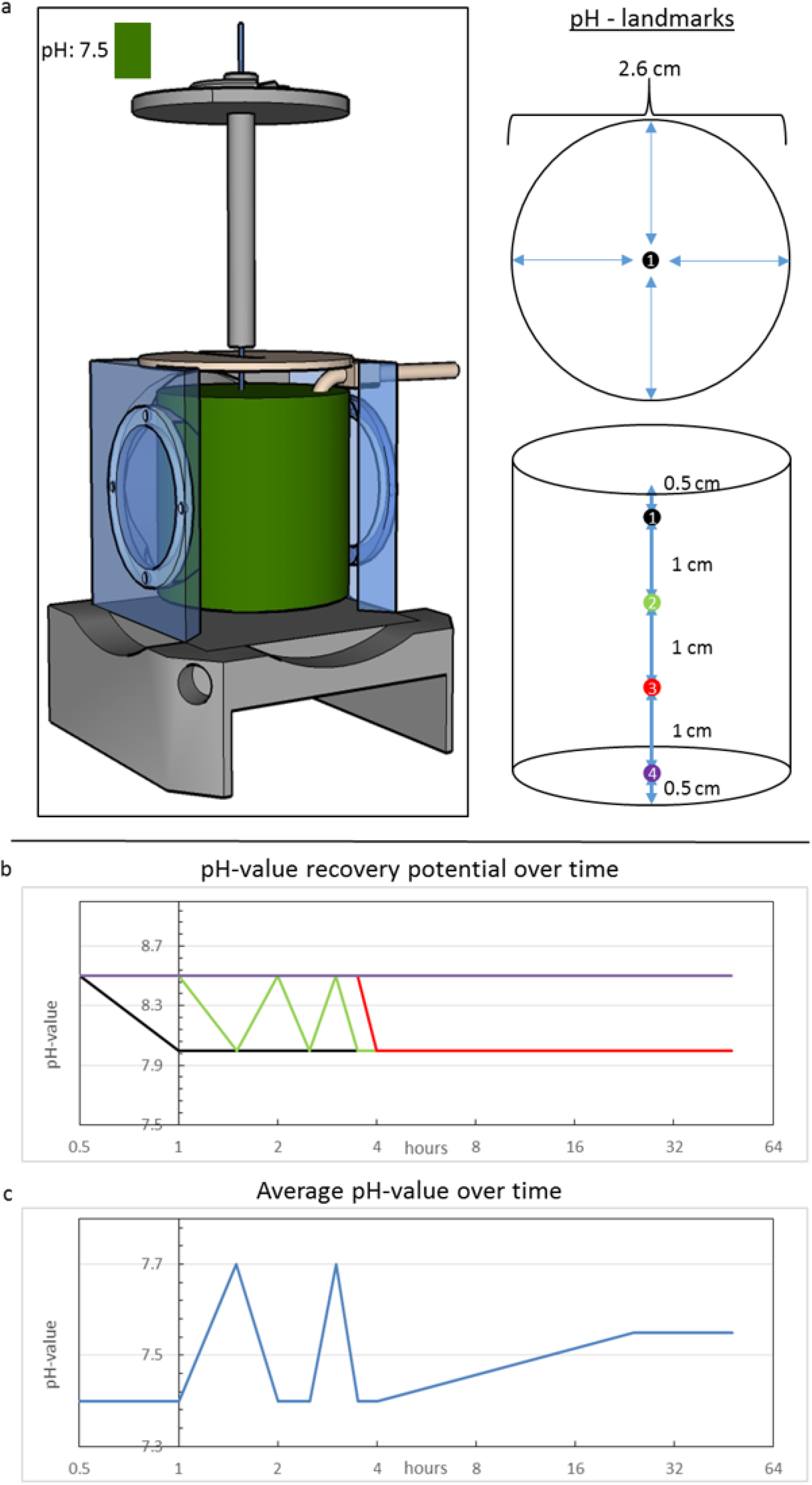
Validation of the pH properties of the Zeiss Lightsheet Z.1 microscope. (**a**) Illustration of the pH-value distribution inside the chamber of the Zeiss Lightsheet Z.1 microscope and the corresponding measurement landmarks. After filling the chamber with buffered media the pH-value is evenly distributed at 7.5 throughout the chamber. (**b**) The constant CO_2_ fumigation that is directed over the liquid column is not able to recover a lower pH-value over time. The pH-value of the medium changes from 8.5 to 8 but it never reaches the physiologically necessary 7.5 (liquid depth: 3 cm). The same is observed at 1 cm and 2 cm liquid depth. At the bottom of the chamber, the pH-value does not change within 48 hours. (**c**) Once the inserted medium has the right pH-value, the incubation system is able to keep it on the same level for more than 2 days.

**Supplementary Figure 5:**
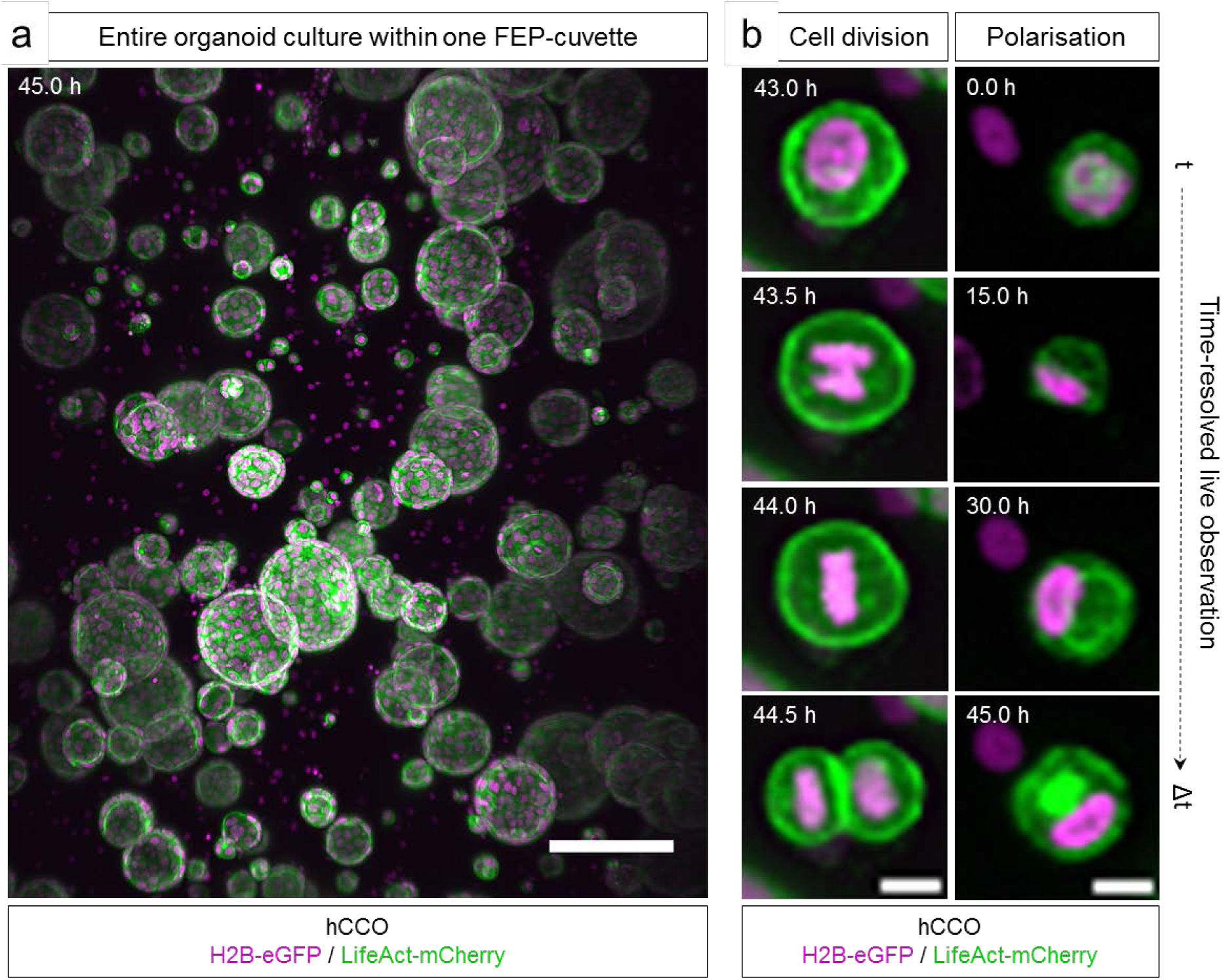
Overview of an entire hCCO culture within one Z1-FEP-cuvette and observation of isolated single-cell dynamics. hCCOs expressed the nuclei marker H2B-eGFP (magenta) and the F-actin cytoskeletal marker LifeAct-mCherry (green). (**a**) Maximum intensity z-projection of the entire field of view in the Lightsheet Z1 microscope. We counted about 120 organoids in this image. Organoids show different sizes and isolated cell nuclei are visible in the interspaces. Scale bar: 50 μm. (**b**) Excerpts of the maximum intensity z-projections shown in (a). Isolated single organoid cells show signs of polarisation and undergo cell division. Microscope: Zeiss Lightsheet Z.1; objective lenses: detection: W Plan-Apochromat 20x/1.0, illumination: Zeiss LSFM 10x/0.2; laser lines: 488 nm, 561 nm; filters: laser block filter (LBF) 405/488/561; voxel size: 1.02 × 1.02 × 2.00 μm^3^; recording interval: 30 min.

**Supplementary Figure 6:**
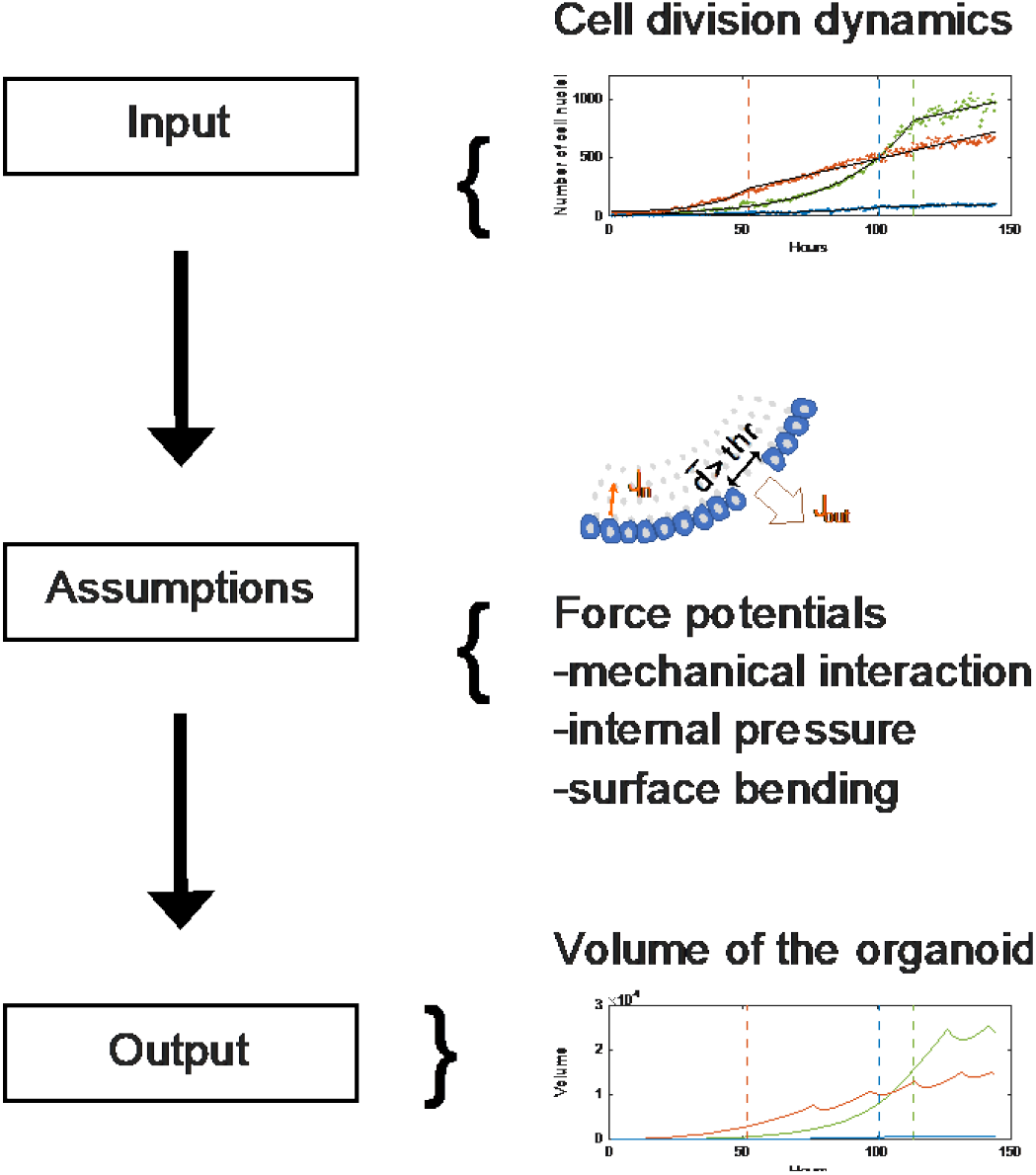
Illustration of the input, the assumptions and the output of the model. Measured cell counts and cell division dynamics are used to initialise the simulations. Organoid behaviour is based on the following main assumptions. (1) Each cell produces a substance with constant rate J_in_, the substance leads to increase of internal pressure. (2) Cell displacement is driven by mechanical cell-cell-interactions, internal pressure and a surface energy of the organoid. (3) If the organoid shell ruptures, substance is released to the outside with flux J_out_, releasing pressure and leading to a contraction of the sphere until the cell-cell connections are restored. The output of the model is the volume data as a function of time of the simulated organoids.

**Supplementary Figure 7:**
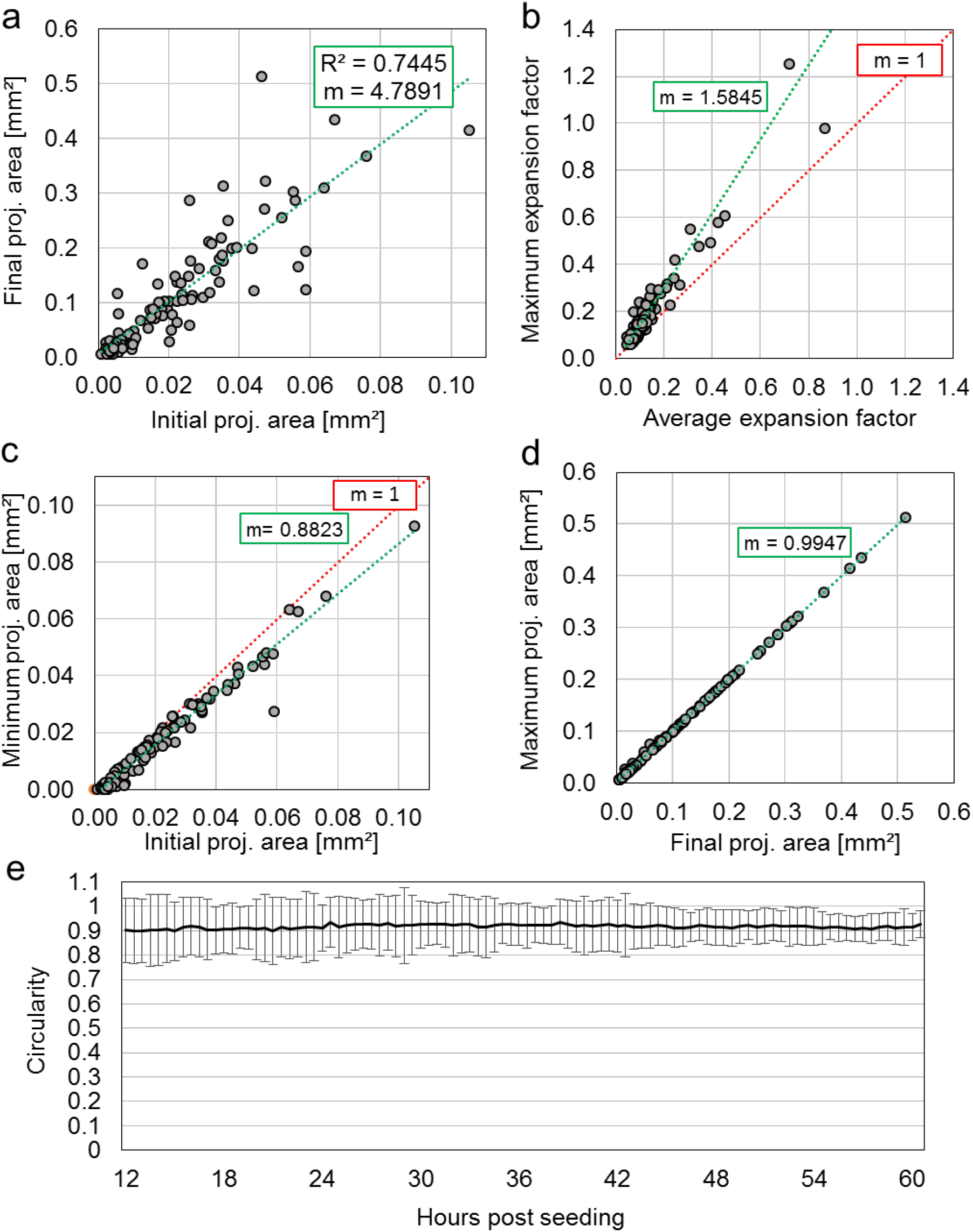
mPO feature extraction using the bright field analysis pipeline. (**a**) The initial and final projected luminal areas correlate positively in healthy mPOs (R^^2^^-value = 0.7445). (**b**) The maximum slope of the expansion phases are in average higher than the average slope. (**c**) The minimum area falls in average slightly below the initial area. (**d**) Furthermore, the final area equals the maximum area, which indicates continuous growth – green: linear trend line, m: slope, red: f(x) = 1x. (**e**) Average circularity over time of organoids grown in three wells. Average standard deviation estimated within the three wells is indicated. Mathematically possible values range between 0 and 1.

**Supplementary Figure 8:**
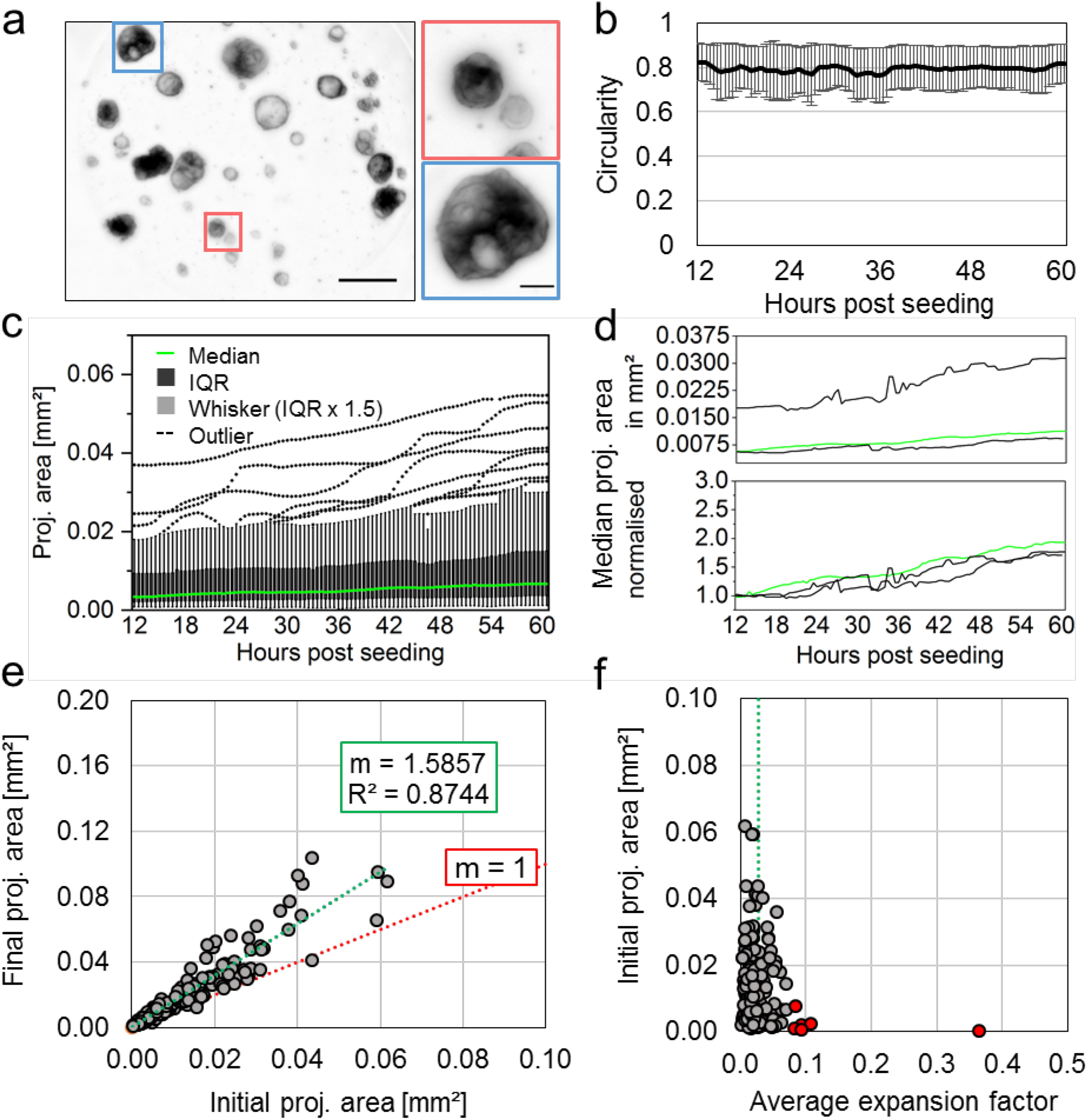
Bright field pipeline allows detailed analysis of polycystic hCCOs. (**a**) Polycystic hCCOs display a dense phenotype. Microscope: Zeiss Axio Observer Z.1; objective lenses: Plan-Apochromat 5x/0.16, avg z-projection, voxel size: 1.29 × 1.29 × 65 μm^^3^^, scale bar overview: 500 μm, close-up: 25 μm. (**b**) The average circularity is around 0.8 over time. (**c**) Similarly to monocystic organoid cultures, the detected projected luminal areas and growth behaviours are heterogeneous. Box plot analysis, median in green (n = 87) (**d**) While the median projected luminal areas of three different wells (technical replicates) vary, the normalised projected area increase is similar (n = 87, 34, 63). (**e**) The initial projected area correlates with the final projected area (R^2^ = 0.8744) with a linear regression slope m = 1.5857. (**f**) The organoids display a similar average expansion factor independent of their initial size with an average of 0.02 and outliers (red) lying above 0.078.

### Supplementary Tables

**Supplementary Table 1:** Used settings for the segmentation and post-processing of the data obtained with the light-sheet pipeline

| Segmentation settings |  | Post processing settings |  |
| --- | --- | --- | --- |
| NucleiFilterRange | 2 | Alpha | 150 |
| NucleiThresholdRange | 10 | OutlierDistanceThreshold | 50 |
| NucleiMeanFactor | 1 | EdgeDistanceThreshold | 50 |
| NucleiStandardDeviationFactor | 0 | NucleiMinCount | 50 |
| NucleiBackgroundFactor | 0.9 | NucleiMaxCount | 10000 |
| HoleFillingRange | 1 |  |  |
| MaxDetectionRange | 0.12 |  |  |
| NucleiSeedDetectionMinRadius | 1 |  |  |
| NucleiSeedDetectionMaxRadius | 4 |  |  |
| NucleiSeedDilation | 1 |  |  |

### Supplementary Movies

**Supplementary Movie 1: Time-resolved observations of epithelial organoids growing in Z1-FEP-cuvettes.** hCCOs expressing H2B-eGFP as nuclei marker (red) and LifeAct-mCherry as F-actin cytoskeletal marker (green) and mPOs expressing Rosa26-nTnG (grey) as nuclei marker were recorded over 10 hours and 143 hours respectively. The formation of organoids from the initially seeded cell clusters, including cell cluster contraction, cell polarisation, lumen formation and expansion can be followed. After about 100 hours of observation some mPOs begin to display signs of degeneration due to extended culturing without further medium exchange. These signs of degeneration include an overall shrinking of the organoid, followed by nuclear condensation and fading of the nuclei signal. Microscope: Zeiss Lightsheet Z.1; detection objective: W Plan-Apochromat 20x/1.0, illumination objective: Zeiss LSFM 10x/0.2; laser lines: 488 nm, 561 nm; filters: laser block filter (LBF) 405/488/561; voxel size: 1.02 × 1.02 × 2.00 μm^**3**^; recording interval: 30 min; scale bar: 50 μm.

**Supplementary Movie 2: Time-resolved 3D volume rendering of the formation process of an entire organoid culture observed within one Z1-FEP-cuvette.** hCCOs expressing H2B-eGFP as nuclei marker (red) and LifeAct-mCherry as F-actin cytoskeletal marker (green) were imaged for 5 days. The movie shows an excerpt of the first 10 hours of the recorded data set. All organoids within the cuvette were segmented and tracked over these first 10 hours of recording. The initial processes of cell cluster contraction, lumen formation and subsequent expansion are shown. Depending on the initial cell-cluster size, organoids differ in the time they need to establish a lumen. Microscope: Zeiss Lightsheet Z.1; detection objective: W Plan-Apochromat 20x/1.0, illumination objective: Zeiss LSFM 10x/0.2; laser lines: 488 nm, 561 nm; filters: laser block filter (LBF) 405/488/561; voxel size: 1.02 × 1.02 × 2.00 μm^**3**^; recording interval: 30 min; 3D rendering and tracking software: Arivis Vision4D.

**Supplementary Movie 3: Time-resolved 3D volume rendering of the fusion process of two organoids.** hCCOs expressing H2B-eGFP as nuclei marker (red) and LifeAct-mCherry as F-actin cytoskeletal marker (green) were recorded in a Z1-FEP-cuvette for 5 days. The movie shows an excerpt of the recorded data of 12 hours spanning the fusion process of two organoids. The fusion process is visualised by 3D volume rendering of the data acquired for the cytoskeletal marker (LifeAct-mCherry – green). After the epithelial monolayers of both organoids touch, they begin to form an opening connecting both lumens within one hour. This opening then expands while cells migrate into one connected monolayer. Microscope: Zeiss Lightsheet Z.1; detection objective: W Plan-Apochromat 20x/1.0, illumination objective: Zeiss LSFM 10x/0.2; laser lines: 488 nm, 561 nm; filters: laser block filter (LBF) 405/488/561; voxel size: 1.02 × 1.02 × 2.00 μm^**3**^; recording interval: 30 min; 3D rendering software: Arivis Vision4D.

**Supplementary Movie 4: Time-resolved 3D volume rendering of intra-organoid luminal dynamics.** hCCOs expressing H2B-eGFP as nuclei marker (red) and LifeAct-mCherry as F-actin cytoskeletal marker (green) were recorded in a Z1-FEP-cuvette for a total of 132 hours. The movie shows data recorded between 84 and 108 hours. Luminal dynamics are visualised by 3D volume rendering of the data acquired for the cytoskeletal marker (LifeAct-mCherry – green) of a large organoid (diameter: ≥ 500 μm), presumably formed after fusion of multiple organoids. We can follow the formation, subsequent retraction and eventual rupture of duct-like structures within the organoid’s lumen. Microscope: Zeiss Lightsheet Z.1; detection objective: W Plan-Apochromat 20x/1.0, illumination objective: Zeiss LSFM 10x/0.2; laser lines: 488 nm, 561 nm; filters: laser block filter (LBF) 405/488/561; voxel size: 1.02 × 1.02 × 2.00 μm^**3**^; recording interval: 30 min; 3D rendering software: Arivis Vision4D.

**Supplementary Movie 5: Time-resolved 3D volume rendering of a growing liver organoid with cell segmentation and tracking.** hCCOs expressing H2B-eGFP as nuclei marker (red) and LifeAct-mCherry as F-actin cytoskeletal marker (green) were recorded in a Z1-FEP-cuvette for a total of 132 hours. The movie shows data recorded between 84 and 108 hours. Red spheres illustrate tracked cell nuclei and rainbow-coloured lines indicate the travelled tracks (colour code: red to blue – timepoint 84 to 108). Single as well as multiple organoid tracking is shown. Rotation as well as uni-directional cell movements are visible. Microscope: Zeiss Lightsheet Z.1; detection objective: W Plan-Apochromat 20x/1.0, illumination objective: Zeiss LSFM 10x/0.2; laser lines: 488 nm, 561 nm; filters: laser block filter (LBF) 405/488/561; voxel size: 1.02 × 1.02 × 2.00 μm^**3**^; recording interval: 30 min; 3D rendering and tracking software: Arivis Vision4D.

**Supplementary Movie 6: Alterations in rotation velocity of neighbouring organoids.** Video shows two mPOs as an excerpt from an entire culture grown within one Z1-FEP-cuvette. The organoids expressed Rosa26-nTnG (grey) as nuclei marker and were imaged over 20 hours. Beside the differences in nuclei size, both organoids showing different behaviour. Cell tracking revealed a rotational motion of the epithelial cell monolayer of the organoid with the small roundish nuclei and no rotational motion of the organoid with the big elongated cell nuclei. The two organoids are in close contact but do not fuse or interact with each other. Microscope: Zeiss Lightsheet Z.1; detection objective: W Plan-Apochromat 20x/1.0, illumination objective: Zeiss LSFM 10x/0.2; laser lines: 488 nm, 561 nm; filters: laser block filter (LBF) 405/488/561; voxel size: 1.02 × 1.02 × 2.00 μm^**3**^; recording interval: 30 min; 3D rendering and tracking software: Arivis Vision4D.

**Supplementary Movie 7: Time-resolved cell tracking during organoid expansion reveals different migration speeds within the epithelial monolayer.** hCCOs expressing H2B-eGFP as nuclei marker (red) and LifeAct-mCherry as F-actin cytoskeletal marker (green) recorded in a Z1-FEP-cuvette for a total of 132 hours. The movie shows data recorded between 84 and 108 hours. Image-based segmentation and 3D-rendering revealed that the cells within an organoid migrate with different speeds during organoid expansion. Cells at the organoid’s “poles” tend to show less or slower migration compared to cells located at the organoid’s equatorial plane (from blue to red: 2-7μm per hour). Microscope: Zeiss Lightsheet Z.1; detection objective: W Plan-Apochromat 20x/1.0, illumination objective: Zeiss LSFM 10x/0.2; laser lines: 488 nm, 561 nm; filters: laser block filter (LBF) 405/488/561; voxel size: 1.02 × 1.02 × 2.00 μm^**3**^; recording interval: 30 min; 3D rendering and tracking software: Arivis Vision4D.

**Supplementary Movie 8: Organoid cell cluster migration prior to organoid formation** Video shows one mPO as an excerpt of an entire culture grown within one Z1-FEP-cuvette. The organoids expressed Rosa26-nTnG (grey) as nuclei marker and were imaged over 20 hours. Prior to organoid formation, the initially seeded organoid cell cluster migrates through the ECM for about 25 hours before the cells rearrange to form a spherical structure. The migrated distance is about 250 μm with an average speed of 10 μm (from green/minimum to red/maximum: 2.5 μm/h – 23 μm/h). Microscope: Zeiss Lightsheet Z.1; detection objective: W Plan-Apochromat 20x/1.0, illumination objective: Zeiss LSFM 10x/0.2; laser lines: 488 nm, 561 nm; filters: laser block filter (LBF) 405/488/561; voxel size: 1.02 × 1.02 × 2.00 μm^**3**^; recording interval: 30 min; 3D rendering and tracking software: Arivis Vision4D.

## Supplementary Theoretical Considerations

### A mechanical model to describe the dynamics of pancreas organoids

To describe the growth and dynamics of a pancreas organoid, we assume it has a roughly spherical shape, with cells forming a monolayer filled with fluid at a different pressure relative to the environment. The volume of the organoid is changed by two mechanisms: a) the influx of liquid caused by an osmotic imbalance or active pumping, and b) cell division. The first mechanism increases the tension between the cells. Cell division, on the other hand, increases the surface of the organoid and reduces tension. If the stress is greater than a critical threshold, at least one cell connection breaks and leakage occur through the organoid shell. The leakage reduces the internal pressure, the monolayer can contract, which in turn allows the ruptured cells to reconnect. Subsequently, the whole process can repeat.

A triangulated-network model was used to simulate the membranes as an elastic surface consisting of cells. Here, the shell of the organoid is described as an infinitely thin elastic surface consisting of hard, spherical beads (the cell’s centres) connected by dynamic bonds to form a triangulated network.

A spring potential acting on neighbouring beads is used to describe the elasticity of the shell and has the typical form

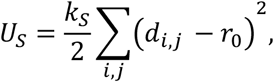

where *k*_*s*_ is the spring constant, *d*_*i,j*_ is the distance between two neighbouring cells *i* and *j*, and *r*_0_ is their equilibrium bond length. The spring potential is minimised if *d*_*i,j*_ between two neighbouring cells *i* and *j* corresponds to the equilibrium distance *r*_0_. Since the method of finite elements is used in the simulation, the force must be derived from the potential used. The force acting on every cell is given as

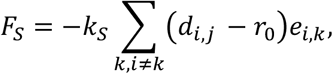

where *k* are the indices of the neighbours of the cell *i*, and *e*_*i,j*_ is the normalised direction vector between ***x***_*i*_ and its neighbour ***x***_*j*_.

The surface bending energy acts on neighbouring triangles and is minimised when the angle between the neighbouring triangles is zero.

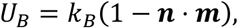

where ***n*** and ***m*** are the normal vectors of two neighbouring triangles sharing a common edge b. The resulting force can be generated by deriving the bending potential after point ***x***_*i*_.

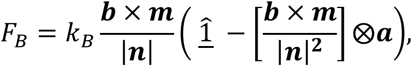

where ***a*** describes the direction vector from ***x***_***i***_ to ***x***_***j***_ in the triangle ***x***_***i***_***x***_***j***_***x***_***k***_ and ***a***⨂***b*** denotes the dynamic product of the two vectors ***a*** and ***b***. For one cell ***x***_***i***_ it is then summed over all ***n***, ***m*** pairs of the neighbouring triangles of ***x***_***i***_. In order to compensate for the differences between cells with different numbers of neighbouring cells, a normalisation is made about the number of neighbouring cells.

In the simulation, the assumption is made that each cell pumps mass (e.g. water) or fluid through an osmotic imbalance into the lumen of the organoid and thus, an internal pressure that differs from the external pressure can build up. The internal pressure is one of the factors of organoid expansion. The force *F*_*P*_, which affects each mass point from the resulting osmotic pressure Π, is given as

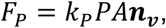

whereby

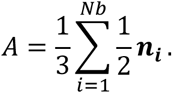

The osmotic pressure Π has an effect on the area *A*, which is understood as the sum of the adjacent triangular areas to the cell center ***x***_***i***_, with the direction vector ***n*** (normal vector to cell *i*= summed normal vectors of the adjacent triangles). The osmotic pressure Π is calculated using the van-‘t-Hoff law for osmotic pressure

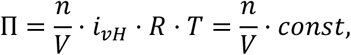

where *n* is the amount of substance, *i*_*vH*_ is the van-‘t-Hoff factor, *R* is the ideal gas constant and *T* is the temperature. The volume of the organoid *V* is calculated for each time step over the convex shell of the cells. The amount of secreted substance changes over time with

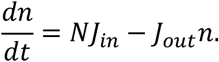

*N* indicates the number of cells in the organoid, *J*_*in*_ the amount of substance produced per cell, and *J*_*out*_ describes the substance drop through a hole in the organoid shell. If the organoid shell has a rupture, *J*_*out*_ is greater than zero, otherwise *J*_*out*_ is zero. The equation of motion used in the simulation applies to the overdamped case and contains stochastic fluctuations *F*_*t*_ of the cells,

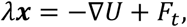

whereby the potential *U* is given as the sum of the above-mentioned potentials.

Cell division is adjusted to the experimental data, obtained by long-term single cell analysis of pancreas-derived organoids, but can easily be adapted to other growth dynamics. If cell division takes place a new cell is added to the system in the middle of a random triangle formed by three neighbouring cells.

## References

Aberle, M. R., Burkhart, R. A., Tiriac, H., Olde Damink, S. W. M., Dejong, C. H. C., Tuveson, D. A., & van Dam, R. M. (2018). Patient-derived organoid models help define personalized management of gastrointestinal cancer. British Journal of Surgery, 105(2), e48–e60. doi: 10.1002/bjs.10726

Bolhaqueiro, A. C. F., Ponsioen, B., Bakker, B., Klaasen, S. J., Kucukkose, E., van Jaarsveld, H., Vivié, J., Verlaan-Klink, I., Hami, N., Spierings, D. C. J., Sasaki, N., Dutta, D., Boj, F., Vries, R. G. J., Lansdorp, P. M., van de Wetering, M., van Oudenaarden, A., Clevers, H., Kranenburg, O.,…Kops, G. J. P. L. (2019). Ongoing chromosomal instability and karyotype evolution in human colorectal cancer organoids. Nature Genetics, 51(5), 824–834. doi: 10.1038/s41588-019-0399-6

Broutier, L., Andersson-Rolf, A., Hindley, C. J., Boj, S. F., Clevers, H., Koo, B.-K., & Huch, M. (2016). Culture and establishment of self-renewing human and mouse adult liver and pancreas 3D organoids and their genetic manipulation. Nature Protocols, 11(9), 1724–1743. doi: 10.1038/nprot.2016.097

Broutier, L., Mastrogiovanni, G., Verstegen, M. M. A., Francies, H. E., Gavarró, L. M., Bradshaw, C. R., Allen, G. E., Arnes-Benito, R., Sidorova, O., Gaspersz, M. P., Georgakopoulos, N., Koo, B. K., Dietmann, S., Davies, S. E., Praseedom, R. K., Lieshout, R., IJzermans, J. N. M., Wigmore, S. J., Saeb-Parsy, K.,…Huch, M. (2017). Human primary liver cancer-derived organoid cultures for disease modeling and drug screening. Nature Medicine, 23(12), 1424–1435. doi: 10.1038/nm.4438

Bruens, L., Ellenbroek, S. I. J., van Rheenen, J., & Snippert, H. J. (2017). In Vivo Imaging Reveals Existence of Crypt Fission and Fusion in Adult Mouse Intestine. Gastroenterology, 153(3), 674–677.e3. doi: 10.1053/j.gastro.2017.05.019

Clevers, H. (2016). Modeling Development and Disease with Organoids. In Cell (Vol. 165, Issue 7, pp. 1586–1597). Cell Press. doi: 10.1016/j.cell.2016.05.082

Dahl-Jensen, S., & Grapin-Botton, A. (2017). The physics of organoids: A biophysical approach to understanding organogenesis. Development (Cambridge), 144(6), 946–951. doi: 10.1242/dev.143693

de Winter-De Groot, K. M., Janssens, H. M., van Uum, R. T., Dekkers, J. F., Berkers, G., Vonk, A., Kruisselbrink, E., Oppelaar, H., Vries, R., Clevers, H., Houwen, R. H. J., Escher, J. C., Elias, S. G., De Jonge, H. R., de Rijke, Y. B., Tiddens, H. A. W. M., van der Ent, C. K., & Beekman, J. M. (2018). Stratifying infants with cystic fibrosis for disease severity using intestinal organoid swelling as a biomarker of CFTR function. European Respiratory Journal, 52(3). doi: 10.1183/13993003.02529-2017

Dossena, M., Piras, R., Cherubini, A., Barilani, M., Dugnani, E., Salanitro, F., Moreth, T., Pampaloni, F., Piemonti, L., & Lazzari, L. (2020). Standardized GMP-compliant scalable production of human pancreas organoids. Stem Cell Research and Therapy, 11(1), 94. doi: 10.1186/s13287-020-1585-2

Dumortier, J. G., Le Verge-Serandour, M., Tortorelli, A. F., Mielke, A., De Plater, L., Turlier, H., & Maître, J. L. (2019). Hydraulic fracturing and active coarsening position the lumen of the mouse blastocyst. Science, 365(6452), 465–468. doi: 10.1126/science.aaw7709

Eils, R., & Kriete, A. (2013). Introducing Computational Systems Biology. In Computational Systems Biology: From Molecular Mechanisms to Disease: Second Edition (pp. 1–8). Elsevier Inc. doi: 10.1016/B978-0-12-405926-9.00001-0

Fan, H., Demirci, U., & Chen, P. (2019). Emerging organoid models: Leaping forward in cancer research. In Journal of Hematology and Oncology (Vol. 12, Issue 1, p. 142). BioMed Central Ltd. doi: 10.1186/s13045-019-0832-4

Fatehullah, A., Tan, S. H., & Barker, N. (2016). Organoids as an in vitro model of human development and disease. In Nature Cell Biology (Vol. 18, Issue 3, pp. 246–254). Nature Publishing Group. doi: 10.1038/ncb3312

Ferrari, A., Veligodskiy, A., Berge, U., Lucas, M. S., & Kroschewski, R. (2008). ROCK-mediated contractility, tight junctions and channels contribute to the conversion of a preapical patch into apical surface during isochoric lumen initiation. Journal of Cell Science, 121(21), 3649–3663. doi: 10.1242/jcs.018648

Georgakopoulos, N., Prior, N., Angres, B., Mastrogiovanni, G., Cagan, A., Harrison, D., Hindley, C. J., Arnes-Benito, R., Liau, S. S., Curd, A., Ivory, N., Simons, B. D., Martincorena, I., Wurst, H., Saeb-Parsy, K., & Huch, M. (2020). Long-term expansion, genomic stability and in vivo safety of adult human pancreas organoids. BMC Developmental Biology, 20(1), 4. doi: 10.1186/s12861-020-0209-5

Greger, K., Swoger, J., & Stelzer, E. H. K. (2007). Basic building units and properties of a fluorescence single plane illumination microscope. Review of Scientific Instruments, 78(2), 023705. doi: 10.1063/1.2428277

Grün, D., Lyubimova, A., Kester, L., Wiebrands, K., Basak, O., Sasaki, N., Clevers, H., & Van Oudenaarden, A. (2015). Single-cell messenger RNA sequencing reveals rare intestinal cell types. Nature, 525(7568), 251–255. doi: 10.1038/nature14966

Harris, T. J. C., & Tepass, U. (2010). Adherens junctions: From molecules to morphogenesis. In Nature Reviews Molecular Cell Biology (Vol. 11, Issue 7, pp. 502–514). Nature Publishing Group. doi: 10.1038/nrm2927

Hirata, E., Ichikawa, T., Horike, S. ichi, & Kiyokawa, E. (2018). Active K-RAS induces the coherent rotation of epithelial cells: A model for collective cell invasion in vitro. Cancer Science, 109(12), 4045–4055. doi: 10.1111/cas.13816

Hötte, K., Koch, M., Hof, L., Tuppi, M., Moreth, T., Verstegen, M. M. A., van der Laan, L. J. W., Stelzer, E. H. K., & Pampaloni, F. (2019). Ultra-thin fluorocarbon foils optimise multiscale imaging of three-dimensional native and optically cleared specimens. Scientific Reports, 9(1), 1–13. doi: 10.1038/s41598-019-53380-2

Huch, M., Bonfanti, P., Boj, S. F., Sato, T., Loomans, C. J. M., Van De Wetering, M., Sojoodi, M., Li, V. S. W., Schuijers, J., Gracanin, A., Ringnalda, F., Begthel, H., Hamer, K., Mulder, J., Van Es, J. H., De Koning, E., Vries, R. G. J., Heimberg, H., & Clevers, H. (2013). Unlimited in vitro expansion of adult bi-potent pancreas progenitors through the Lgr5/R-spondin axis. EMBO Journal, 32(20), 2708–2721. doi: 10.1038/emboj.2013.204

Huch, M., Gehart, H., van Boxtel, R., Hamer, K., Blokzijl, F., Verstegen, M. M. A., Ellis, E., van Wenum, M., Fuchs, S. A., de Ligt, J., van de Wetering, M., Sasaki, N., Boers, S. J., Kemperman, H., de Jonge, J., Ijzermans, J. N. M., Nieuwenhuis, E. E. S., Hoekstra, R., Strom, S.,…Clevers, H. (2015). Long-Term Culture of Genome-Stable Bipotent Stem Cells from Adult Human Liver. Cell, 160(1–2), 299–312. doi: 10.1016/J.CELL.2014.11.050

Huch, M., Knoblich, J. A., Lutolf, M. P., & Martinez-Arias, A. (2017). The hope and the hype of organoid research. Development (Cambridge), 144(6), 938–941. doi: 10.1242/dev.150201

Ishiguro, H., Yamamoto, A., Nakakuki, M., Yi, L., Ishiguro, M., Yamaguchi, M., Kondo, S., & Mochimaru, Y. (2012). Physiology and pathophysiology of bicarbonate secretion by pancreatic duct epithelium. In Nagoya J. Med. Sci (Vol. 74, Issue 1).

Karolak, A., Markov, D. A., McCawley, L. J., & Rejniak, K. A. (2018). Towards personalized computational oncology: From spatial models of tumour spheroids, to organoids, to tissues. In Journal of the Royal Society Interface (Vol. 15, Issue 138). Royal Society Publishing. doi: 10.1098/rsif.2017.0703

Keller, P. J., Schmidt, A. D., Wittbrodt, J., & Stelzer, E. H. K. (2008). Reconstruction of zebrafish early embryonic development by scanned light sheet microscopy. Science, 322(5904), 1065–1069. doi: 10.1126/science.1162493

Kim, S., Lewis, A. E., Singh, V., Ma, X., Adelstein, R., & Bush, J. O. (2015). Convergence and Extrusion Are Required for Normal Fusion of the Mammalian Secondary Palate. PLoS Biology, 13(4), 1–25. doi: 10.1371/journal.pbio.1002122

Kretzschmar, K., & Clevers, H. (2016). Organoids: Modeling Development and the Stem Cell Niche in a Dish. In Developmental Cell (Vol. 38, Issue 6, pp. 590–600). Cell Press. doi: 10.1016/j.devcel.2016.08.014

Lancaster, M. A., & Huch, M. (2019). Disease modelling in human organoids. DMM Disease Models and Mechanisms, 12(7), dmm039347. doi: 10.1242/dmm.039347

Lancaster, M. A., Renner, M., Martin, C. A., Wenzel, D., Bicknell, L. S., Hurles, M. E., Homfray, T., Penninger, J. M., Jackson, A. P., & Knoblich, J. A. (2013). Cerebral organoids model human brain development and microcephaly. Nature, 501(7467), 373–379. doi: 10.1038/nature12517

Legland, D., Arganda-Carreras, I., & Andrey, P. (2016). MorphoLibJ: Integrated library and plugins for mathematical morphology with ImageJ. Bioinformatics, 32(22), 3532–3534. doi: 10.1093/bioinformatics/btw413

Loomans, C. J. M., Williams Giuliani, N., Balak, J., Ringnalda, F., van Gurp, L., Huch, M., Boj, S. F., Sato, T., Kester, L., de Sousa Lopes, S. M. C., Roost, M. S., Bonner-Weir, S., Engelse, M. A., Rabelink, T. J., Heimberg, H., Vries, R. G. J., van Oudenaarden, A., Carlotti, F., Clevers, H., & de Koning, E. J. P. (2018). Expansion of Adult Human Pancreatic Tissue Yields Organoids Harboring Progenitor Cells with Endocrine Differentiation Potential. Stem Cell Reports, 10(3), 712–724. doi: 10.1016/j.stemcr.2018.02.005

Mahe, M. M., Aihara, E., Schumacher, M. A., Zavros, Y., Montrose, M. H., Helmrath, M. A., Sato, T., & Shroyer, N. F. (2013). Establishment of Gastrointestinal Epithelial Organoids. Current Protocols in Mouse Biology, 3(4), 217–240. doi: 10.1002/9780470942390.mo130179

Mandelkow, R., GüMBEL, D., Ahrend, H., Kaul, A., Zimmermann, U., Burchardt, M., & Stope, M. B. (2017). Detection and quantification of nuclear morphology changes in apoptotic cells by fluorescence microscopy and subsequent analysis of visualized fluorescent signals. Anticancer Research, 37(5), 2239–2244. doi: 10.21873/anticanres.11560

Marmaras, A., Berge, U., Ferrari, A., Kurtcuoglu, V., Poulikakos, D., & Kroschewski, R. (2010). A mathematical method for the 3D analysis of rotating deformable systems applied on lumen-forming MDCK cell aggregates. Cytoskeleton, 67(4), 224–240. doi: 10.1002/cm.20438

Montes-Olivas, S., Marucci, L., & Homer, M. (2019). Mathematical Models of Organoid Cultures. Frontiers in Genetics, 10, 873. doi: 10.3389/fgene.2019.00873

Nagle, P. W., Plukker, J. T. M., Muijs, C. T., van Luijk, P., & Coppes, R. P. (2018). Patient-derived tumor organoids for prediction of cancer treatment response. In Seminars in Cancer Biology (Vol. 53, pp. 258–264). Academic Press. doi: 10.1016/j.semcancer.2018.06.005

Odenwald, M. A., Choi, W., Buckley, A., Shashikanth, N., Joseph, N. E., Wang, Y., Warren, M. H., Buschmann, M. M., Pavlyuk, R., Hildebrand, J., Margolis, B., Fanning, A. S., & Turner, J. R. (2017). ZO-1 interactions with F-actin and occludin direct epithelial polarization and single lumen specification in 3D culture. Journal of Cell Science, 130(1), 243–259. doi: 10.1242/jcs.188185

Ooft, S. N., Weeber, F., Dijkstra, K. K., McLean, C. M., Kaing, S., van Werkhoven, E., Schipper, L., Hoes, L., Vis, D. J., van de Haar, J., Prevoo, W., Snaebjornsson, P., van der Velden, D., Klein, M., Chalabi, M., Boot, H., van Leerdam, M., Bloemendal, H. J., Beerepoot, L. V.,…Voest, E. E. (2019). Patient-derived organoids can predict response to chemotherapy in metastatic colorectal cancer patients. Science Translational Medicine, 11(513). doi: 10.1126/scitranslmed.aay2574

Preibisch, S., Saalfeld, S., & Tomancak, P. (2009). Globally optimal stitching of tiled 3D microscopic image acquisitions. Bioinformatics, 25(11), 1463–1465. doi: 10.1093/bioinformatics/btp184

Ruiz-Herrero, T., Alessandri, K., Gurchenkov, B. V., Nassoy, P., & Mahadevan, L. (2017). Organ size control via hydraulically gated oscillations. Development, 144(23), 4422–4427. doi: 10.1242/dev.153056

Sasai, Y. (2013). Cytosystems dynamics in self-organization of tissue architecture. Nature, 493(7432), 318–326. doi: 10.1038/nature11859

Schlaermann, P., Toelle, B., Berger, H., Schmidt, S. C., Glanemann, M., Ordemann, J., Bartfeld, S., Mollenkopf, H. J., & Meyer, T. F. (2016). A novel human gastric primary cell culture system for modelling Helicobacter pylori infection in vitro. Gut, 65(2), 202–213. doi: 10.1136/gutjnl-2014-307949

Schmitz, A., Fischer, S. C., Mattheyer, C., Pampaloni, F., & Stelzer, E. H. K. (2017). Multiscale image analysis reveals structural heterogeneity of the cell microenvironment in homotypic spheroids. Scientific Reports, 7(January), 43693 (1-13). doi: 10.1038/srep43693

Schwank, G., Andersson-Rolf, A., Koo, B. K., Sasaki, N., & Clevers, H. (2013). Generation of BAC Transgenic Epithelial Organoids. PLoS ONE, 8(10), 6–11. doi: 10.1371/journal.pone.0076871

Sebrell, T. A., Sidar, B., Bruns, R., Wilkinson, R. A., Wiedenheft, B., Taylor, P. J., Perrino, B. A., Samuelson, L. C., Wilking, J. N., & Bimczok, D. (2018). Live imaging analysis of human gastric epithelial spheroids reveals spontaneous rupture, rotation and fusion events. Cell and Tissue Research, 371(2), 293–307. doi: 10.1007/s00441-017-2726-5

Serra, D., Mayr, U., Boni, A., Lukonin, I., Rempfler, M., Challet Meylan, L., Stadler, M. B., Strnad, P., Papasaikas, P., Vischi, D., Waldt, A., Roma, G., & Liberali, P. (2019). Self-organization and symmetry breaking in intestinal organoid development. Nature, 569, 66–72. doi: 10.1038/s41586-019-1146-y

Stelzer, E. H. K. (2015). Light-sheet fluorescence microscopy for quantitative biology. Nature Methods, 12(1), 23–26. doi: 10.1038/nmeth.3219

Stichel, D., Middleton, A. M., Müller, B. F., Depner, S., Klingmüller, U., Breuhahn, K., & Matthäus, F. (2017). An individual-based model for collective cancer cell migration explains speed dynamics and phenotype variability in response to growth factors. Npj Systems Biology and Applications, 3(1), 1–10. doi: 10.1038/s41540-017-0006-3

Takeda, N., Jain, R., Li, D., Li, L., Lu, M. M., & Epstein, J. A. (2013). Lgr5 Identifies Progenitor Cells Capable of Taste Bud Regeneration after Injury. PLoS ONE, 8(6), e66314. doi: 10.1371/journal.pone.0066314

Tanner, K., Mori, H., Mroue, R., Bruni-Cardoso, A., & Bissell, M. J. (2012). Coherent angular motion in the establishment of multicellular architecture of glandular tissues. Proceedings of the National Academy of Sciences of the United States of America, 109(6), 1973–1978. doi: 10.1073/pnas.1119578109

Trepat, X., & Sahai, E. (2018). Mesoscale physical principles of collective cell organization. In Nature Physics (Vol. 14, Issue 7, pp. 671–682). Nature Publishing Group. doi: 10.1038/s41567-018-0194-9

Verveer, P. J., Swoger, J., Pampaloni, F., Greger, K., Marcello, M., & Stelzer, E. H. K. (2007). High-resolution three-dimensional imaging of large specimens with light sheet–based microscopy. Nature Methods, 4(4), 311–313. doi: 10.1038/nmeth1017

Wang, H., Lacoche, S., Huang, L., Xue, B., & Muthuswamy, S. K. (2013). Rotational motion during three-dimensional morphogenesis of mammary epithelial acini relates to laminin matrix assembly. Proceedings of the National Academy of Sciences of the United States of America, 110(1), 163–168. doi: 10.1073/pnas.1201141110

Xavier da Silveira dos Santos, A., & Liberali, P. (2019). From single cells to tissue self‐ organization. The FEBS Journal, 286(8), 1495–1513. doi: 10.1111/febs.14694

Yang, Q., Xue, S.-L., Chan, C. J., Rempfler, M., Vischi, D., Gutierrez, F. M., Hiiragi, T., Hannezo, E., & Liberali, P. (2020). Cell fate coordinates mechano-osmotic forces in intestinal crypt morphogenesis. BioRxiv, 2020.05.13.094359. doi: 10.1101/2020.05.13.094359

