## Supplementary figures and images for "Long-term live imaging of epithelial organoids and corresponding multiscale analysis reveal high heterogeneity and identify core regulatory principles"

### Supplemental Figure 1

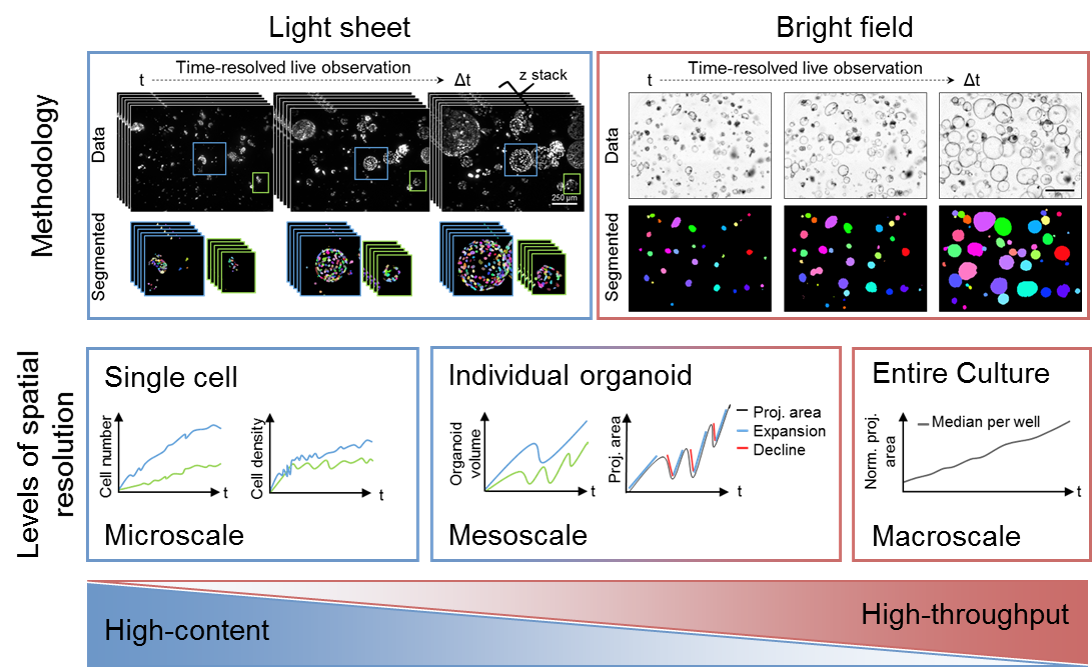

### Supplemental Figure 2

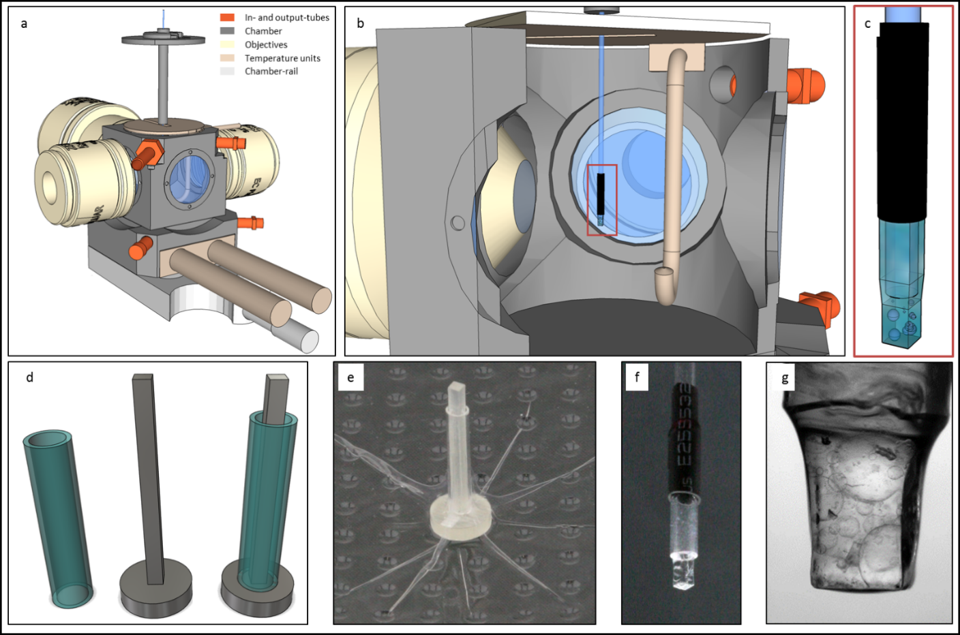

### Supplemental Figure 3

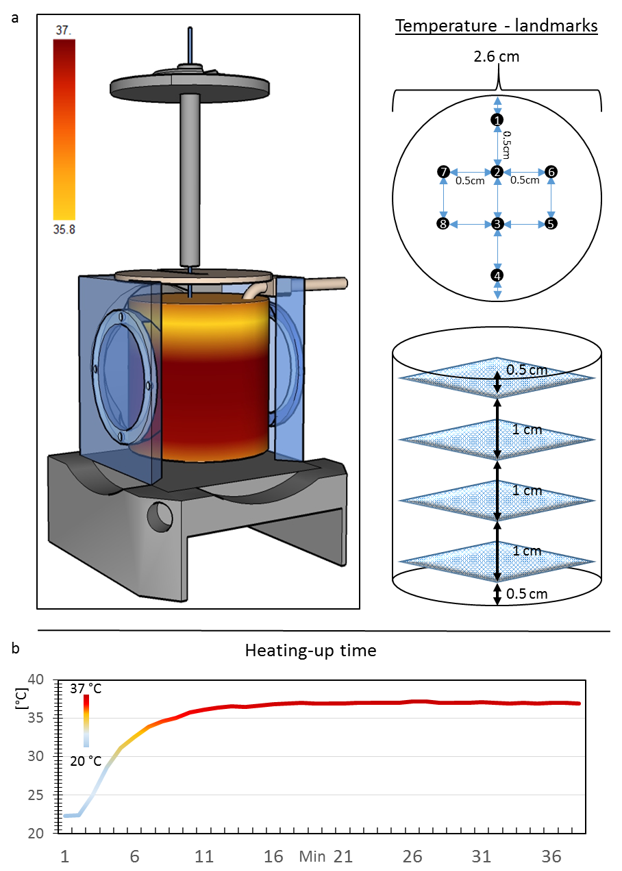

### Supplemental Figure 4

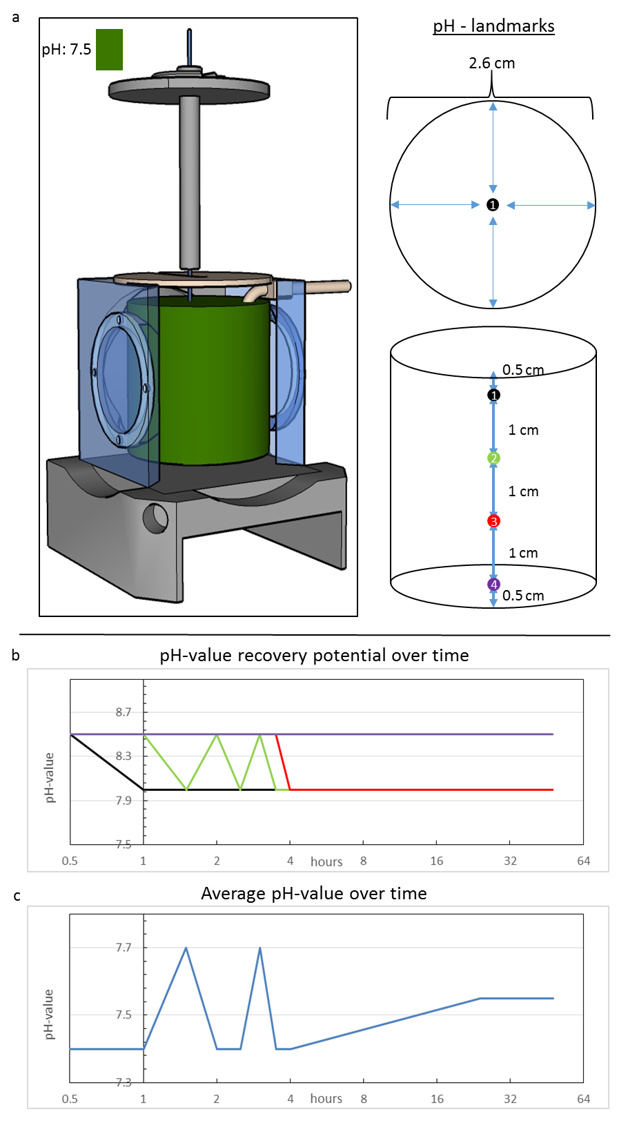

### Supplemental Figure 5

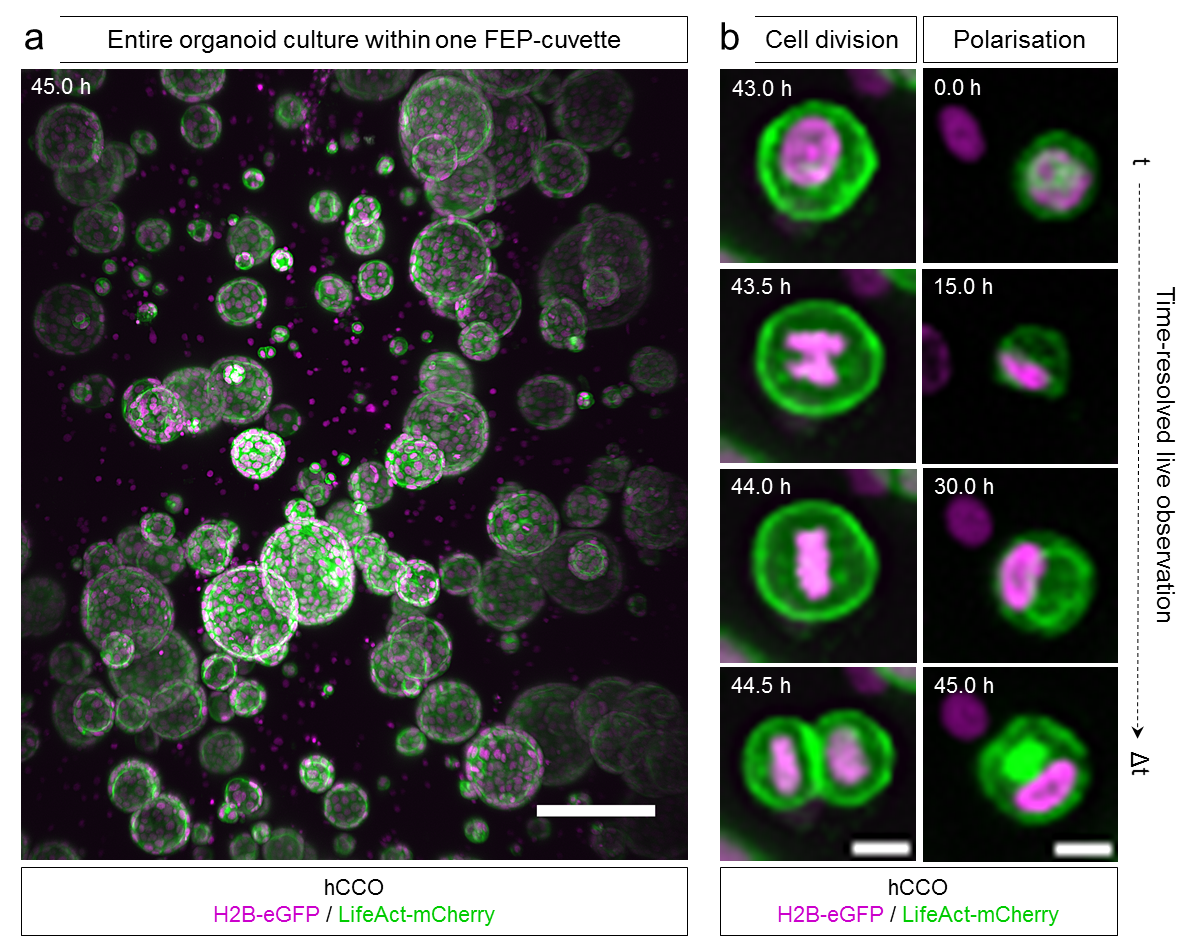

### Supplemental Figure 6

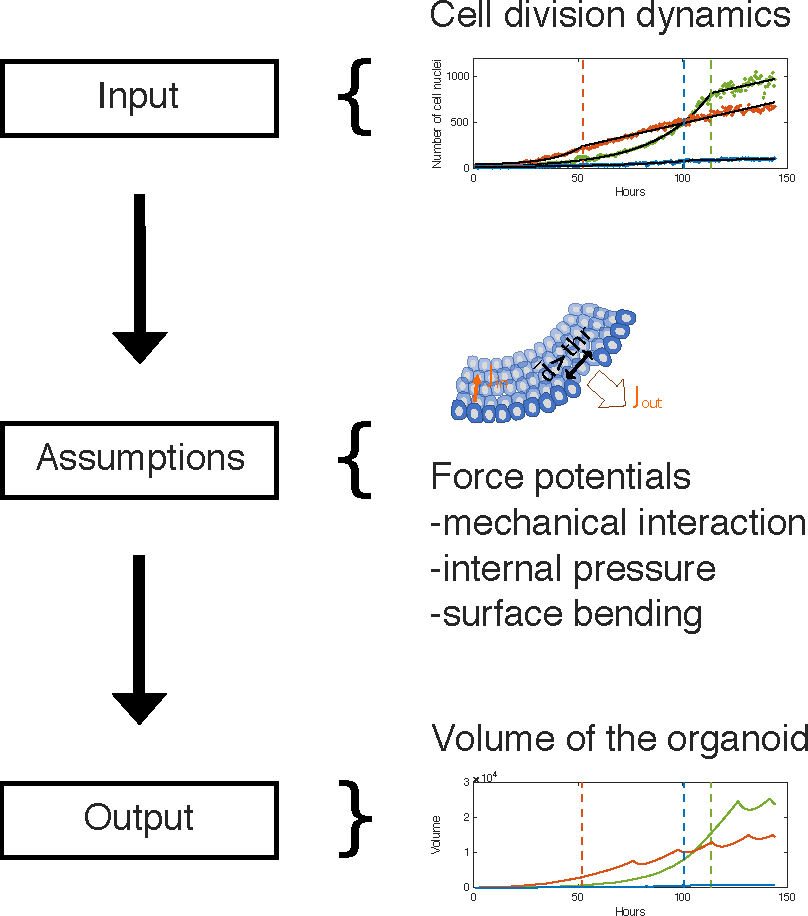

### Supplemental Figure 7

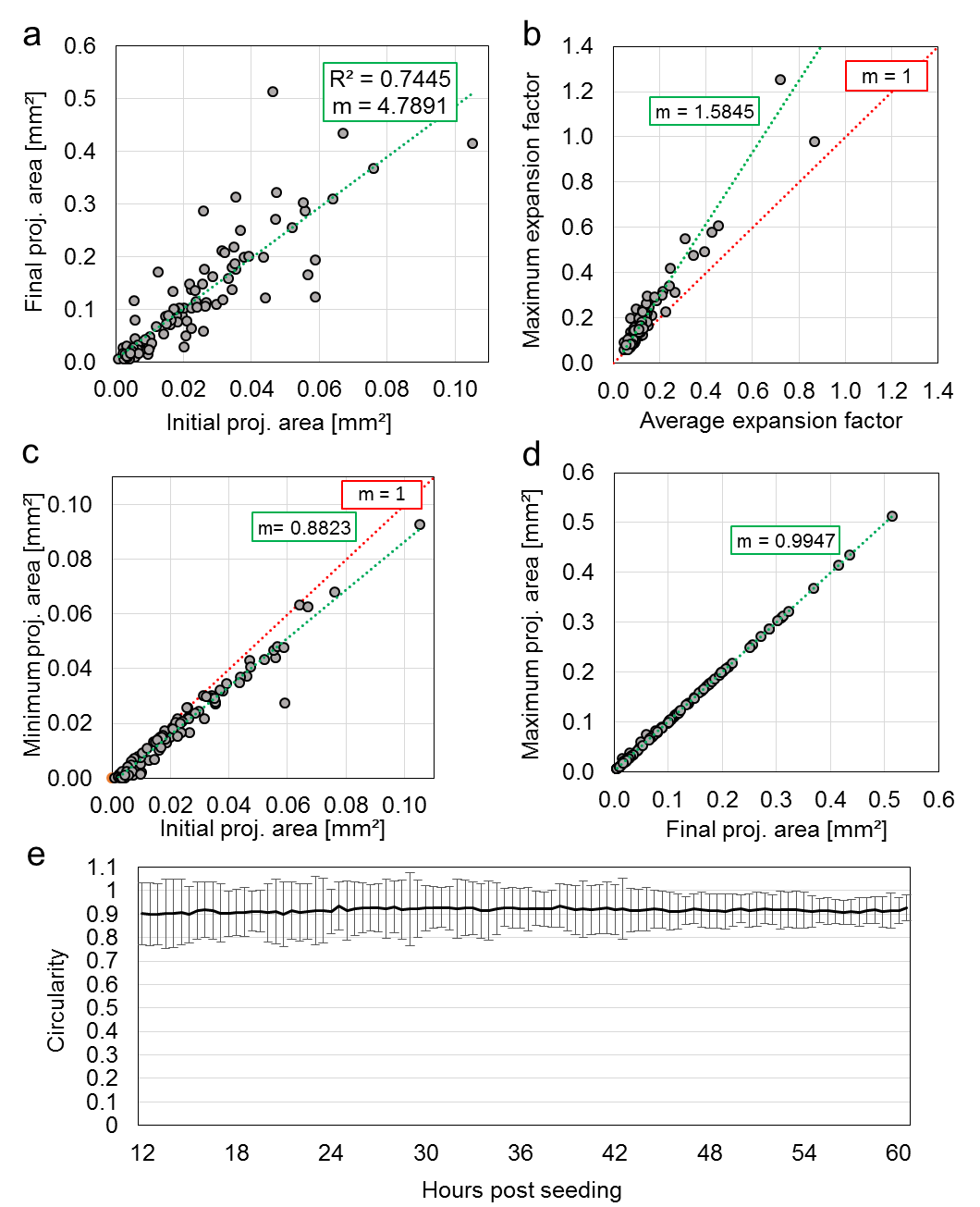

### Supplemental Figure 8

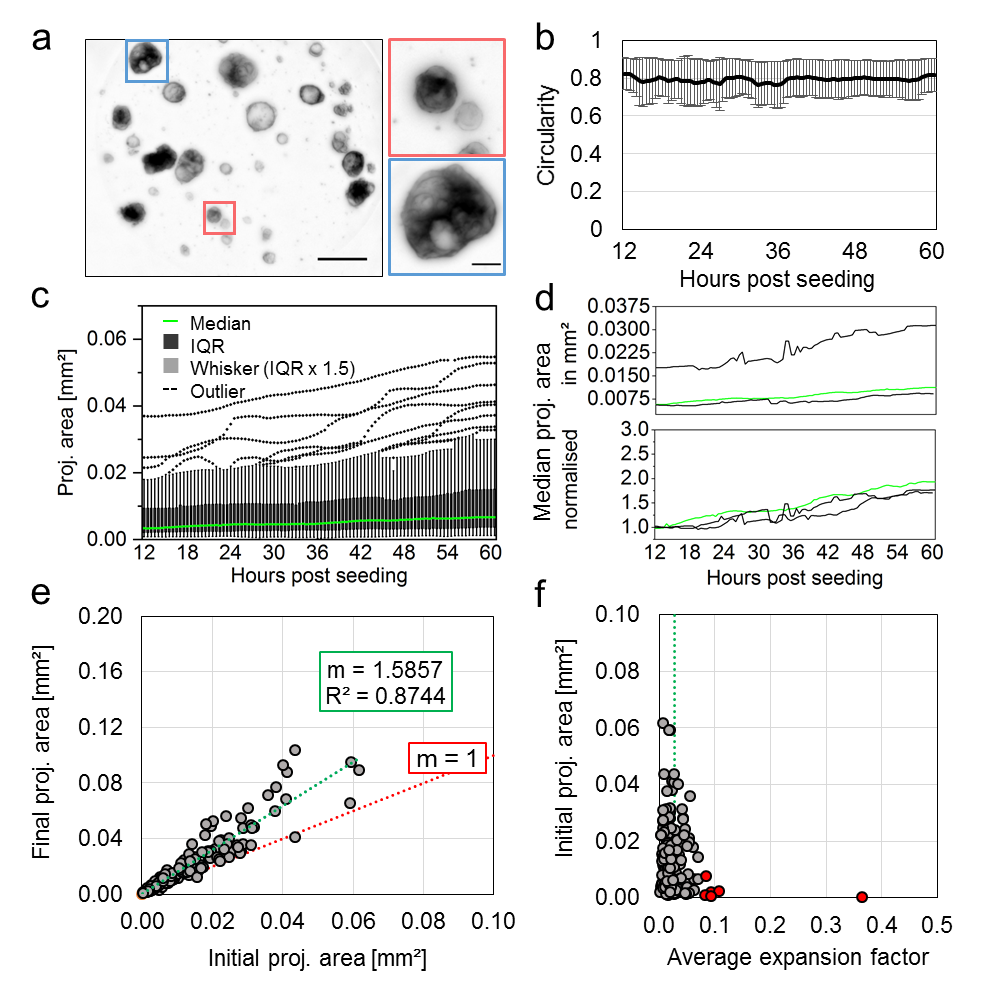
